# Reciprocal regulation among TRPV1 channels and phosphoi nos itide 3-kinase in response to nerve growth factor

**DOI:** 10.1101/346718

**Authors:** Anastasiia Stratiievska, Sara Nelson, Eric N. Senning, Jonathan D. Lautz, Stephen E.P. Smith, Sharona E. Gordon

## Abstract

Although it has been known for over a decade that the inflammatory mediator NGF sensitizes pain-receptor neurons through increased trafficking of TRPV1 channels to the plasma membrane, the mechanism by which this occurs remains mysterious. NGF activates phosphoinositide 3-kinase (PI3K), the enzyme that generates PIP_3_, and PI3K activity is required for sensitization. One tantalizing hint came from the finding that the N-terminal region of TRPV1 interacts directly with PI3K. Using 2-color total internal reflection fluorescence microscopy, we show that TRPV1 potentiates NGF-induced PI3K activity. A soluble TRPV1 fragment corresponding to the N-terminal Ankyrin repeats domain (ARD) was sufficient to produce this potentiation, indicating that allosteric regulation was involved. Further, other TRPV channels with conserved ARDs also potentiated NGF-induced PI3K activity whereas TRP channels lacking ARDs did not. Our data demonstrate a novel reciprocal regulation of PI3K signaling by the ARD of TRPV channels.

## Introduction

Although the current opioid epidemic highlights the need for improved pain therapies, in particular for pain in chronic inflammation (Johannes et al., 2010). Too little is known about the mechanisms that mediate increased sensitivity to pain that occurs in the setting of injury and inflammation (Ji et al., 2014). Inflammatory hyperalgesia, the hypersensitivity to thermal, chemical, and mechanical stimuli (Cesare and McNaughton, 1996), can be divided in two phases, acute and chronic (Dickenson and Sullivan, 1987). Locally released inflammatory mediators, for example, growth factors, bradykinin, prostaglandins, ATP and tissue acidification, (Kozik et al., 1998, Lardner, 2001, Tissot et al., 1989, Burnstock, 1972), directly stimulate and sensitize nociceptive fibers of primary sensory neurons (Cesare and McNaughton, 1996, Bevan and Yeats, 1991, Trebino et al., 2003, Hamilton et al., 1999, McMahon et al., 1995).

One of the proteins that has been studied for its role in hyperalgesia is Transient Receptor Potential Vanilloid Subtype 1 (TRPV1). TRPV1 is a non-selective cation channel that is activated by a variety of noxious stimuli including heat, extracellular protons, and chemicals including capsaicin, a spicy compound in chili pepper (Caterina et al., 1999). TRPV1 is expressed in sensory nociceptive neurons, which are characterized by cell bodies located in the dorsal root ganglia (DRG) and trigeminal ganglia (Caterina et al., 1999). Sensory afferents from these neurons project to skin and internal organs, and synapse onto interneurons in the dorsal horn of the spinal cord (Willis WJ, 1978). TRPV1 activation leads to calcium influx, which results in action potential generation in the sensory neuron and, ultimately, pain sensation (Caterina et al., 1997).

The importance of TRPV1 in inflammatory hyperalgesia was demonstrated by findings that the TRPV1 knock-out mouse showed decreased thermal pain responses and impaired inflammation-induced hyperalgesia (Caterina et al., 2000). TRPV1 activity is enhanced during inflammation which leads to increased pain and lowered pain thresholds (Davis et al., 2000, Zhang et al., 2005, Shu and Mendell, 1999). TRPV1 is modulated by G-protein coupled receptors (GPCRs) and Receptor tyrosine kinases (RTKs), but the mechanism by which these receptors modulate and sensitize TRPV1 is controversial (Suh and Oh, 2005, Shu and Mendell, 1999, Cesare and McNaughton, 1996).

Nerve growth factor (NGF) is one of the best studied RTK agonists involved in inflammatory hyperalgesia (Vetter et al., 1991). NGF acts directly on peptidergic C-fiber nociceptors (Donnerer et al., 1992), which express RTK receptors for NGF: Tropomyosin-receptor-kinase A (TrkA) (McMahon et al., 1995) and neurotrophin receptor p75_NTR_ (Lee et al., 1992). NGF binding to TrkA/p75_NTR_ induces receptor auto-phosphorylation and activation of downstream signaling pathways including phospholipase C (PLC), mitogen-activated protein kinase (MAPK), and phosphoinositide 3-kinase (PI3K) (Vetter et al., 1991, Raffioni and Bradshaw, 1992, Dikic et al., 1995). We and others have previously shown that the acute phase of NGF-induced sensitization requires activation of PI3K, which increases trafficking of TRPV1 channels to the PM (Stein et al., 2006, Bonnington and McNaughton, 2003). In chronic pain, NGF also produces changes in the protein expression of ion channels such as TRPV1 and NaV1.8 (Ji et al., 2002, Thakor et al., 2009, Keh et al., 2008). The acute and chronic phases of the NGF response result in increased “gain” to painful stimuli.

PI3K is a lipid kinase, which phosphorylates the signaling lipid Phosphatidylinositol 4,5-bisphosphate (PIP_2_) into Phosphatidylinositol (3,4,5)-trisphosphate (PIP_3_) (Cantley et al., 1991). PIP_3_ is a signaling lipid as well, and its role in membrane trafficking is well-established (Insall and Weiner, 2001). PI3K is an obligatory heterodimer that includes catalytic p110 and regulatory p85 subunits (Hiles et al., 1992, Hu et al., 1993). The p85 subunit contains two Src homology 2 (SH2) domains (Escobedo et al., 1991), which recognize the phospho-tyrosine motif Y-X-X-M of many activated RTKs and adaptor proteins (Songyang et al., 1993). In the resting state, p85 inhibits the enzymatic activity of p110 via one of its SH2 domains (Miled et al., 2007). This autoinhibition is relieved when p85 binds to a phosphotyrosine motif (Miled et al., 2007). NGF-induced PI3K activity leads to an increase in the number of TRPV1 channels at the PM (Figure 1) (Bonnington and McNaughton, 2003, Stein et al., 2006).

We have previously shown that TRPV1 and p85 interact directly (Stein et al., 2006). We localized the TRPV1/p85 interaction to the N-terminal region of TRPV1 and a region including two SH2 domains of p85 (Stein et al., 2006). However, whether the TRPV1/p85 interaction contributes to NGF-induced trafficking of TRPV1 is unknown. Here, we further localized the functional interaction site for p85 to the region of TRPV1 N-terminus containing several conserved Ankyrin repeats (here we refer to it as Ankyrin repeat domain (ARD)). Remarkably, we found that TRPV1 potentiated the activity of PI3K and that a soluble TRPV1 fragment corresponding to the ARD was sufficient for this potentiation. Because the ARD is structurally conserved among TRPV channels, we tested whether other TRPV channels could also potentiate NGF-induced PI3K activity. We found that TRPV2 and TRPV4 both potentiated NGF-induced PI3K activity and trafficked to the PM in response to NGF. Interestingly, TRPA1 was also trafficked to the plasma membrane in response to NGF, although potentiation of NGF-induced PI3K activity did not achieve statistical significance. In contrast, this reciprocal regulation was not observed for non-ARD containing channels TRPM4 and TRPM8. Together, our data reveal a previously unknown reciprocal regulation among TRPV channels and PI3K. We speculate that this reciprocal regulation could be important wherever TRPV channels are co-expressed with PI3K-coupled RTKs.

**Figure 1.**
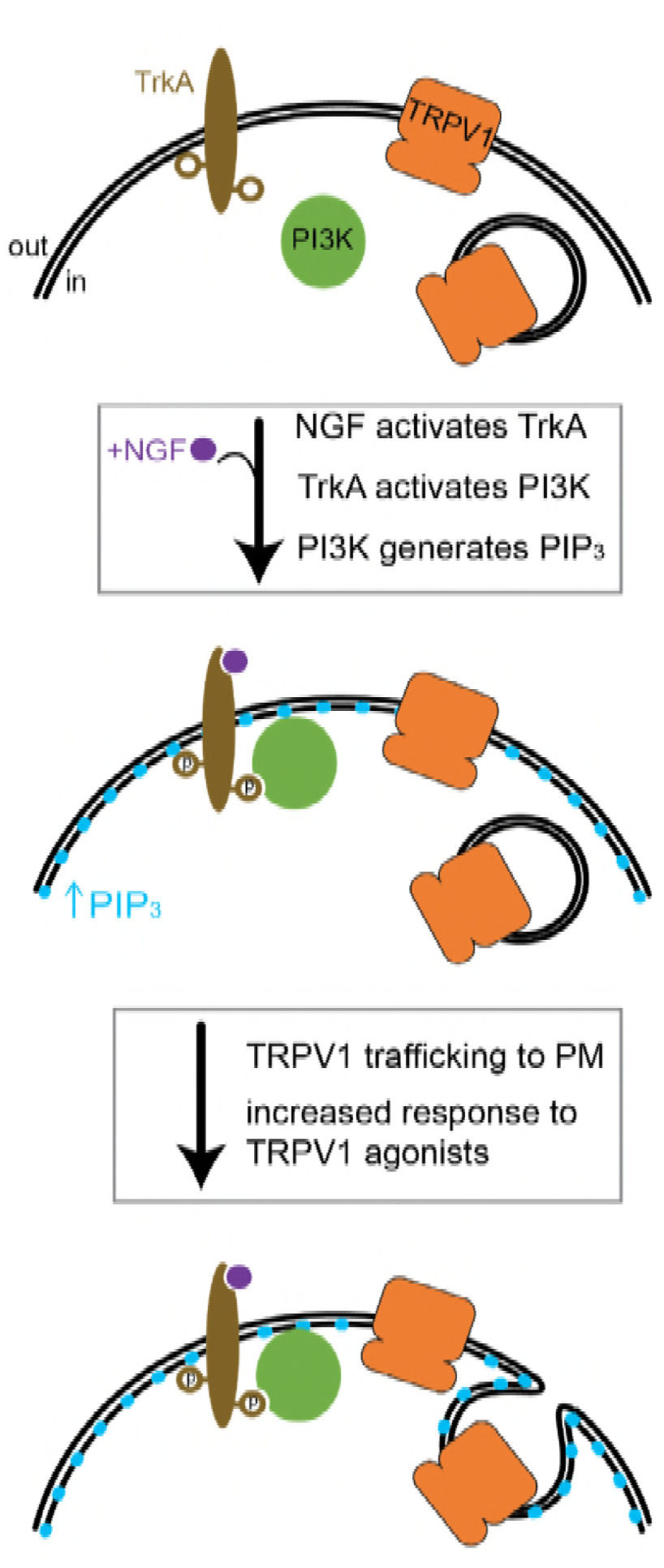
Cartoon depicting the signaling pathway leading to TRPV1 sensitization by NGF. Upon NGF binding, TrkA/p75_NTR_ (labeled as TrkA for simplicity) auto-phosphorylation leads to activation of PI3K and PIP_3_ synthesis. Signaling lipid PIP_3_ leads to signaling events resulting in exocytosis of TRPV1-bearing vesicles and therefore increase in channel number on the plasma membrane.

## Methods

### TIRF Microscopy and analysis

For imaging, we used an inverted microscope (NIKON Ti-E) equipped for total internal fluorescence (TIRF) imaging with a 60x objective (NA 1.49). Glass coverslips with adherent cells were placed in a custom-made chamber. The chamber volume (^~^1 ml) was exchanged using a gravity-driven perfusion system. Cells were acclimated to flow for at least 15 min prior to NGF application. Akt-PH fused to Cyan Fluorescent Protein (CFP) was imaged using excitation from a 447 nm laser and a 480/40 emission filter. TRPV1 fused to Yellow Fluorescent Protein (YFP) was imaged using the 514 nm line of an argon laser and a 530 long pass emission filter. Time-lapse images were obtained by taking consecutive CFP and YFP images every 10 seconds. Movies were then processed using ImageJ software (NIH) (Rasband, 1997-2016). Regions of interest (ROI) were drawn around the footprint of individual cells and the average ROI pixel intensity was measured. Measurements were analyzed using Excel 2013 (Microsoft Corporation), by subtracting the background ROI intensity from the intensity of each cell ROI. Traces were normalized by the average intensity during the 1 min time period prior to NGF application.

### Depth of TIRF field and membrane translocation estimation

Because PIP_3_ levels reported by the Akt-PH fluorescence measured with TIRF microscopy include significant contamination from free Akt-PH in the cytosol, we used the characteristic decay of TIRF illumination to estimate the fraction of our signal due to Akt-PH bound to the membrane. We first estimated the fraction of the illumination at the membrane in resting cells, assuming that free Akt-PH is homogeneously distributed throughout the evanescent field. After stimulation with NGF, we then used this fraction of illumination at the membrane to determine the fraction of the emission light originating from this region. The estimation approach used below was not used to quantitatively evaluate our data. Rather, it demonstrates the general issue of cytosolic contamination causing underestimation of changes in membrane-associated fluorescence even when using TIRF microscopy.

The depth of the TIRF field was estimated as described in the literature (Axelrod, 1981, Mattheyses and Axelrod, 2006). Briefly, when laser light goes through the interface between a coverslip with refractive index *n_2_* and saline solution with refractive index *n_1_*, it experiences total internal reflection at angles less than the critical incidence angle, θ_c_, given by

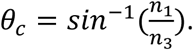

The characteristic depth of the illuminated field *d* is described by

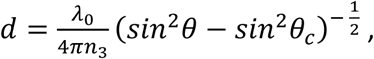

where *λ_0_* is laser wavelength. The illumination decay *τ*, depends on depth of field as follows:

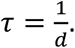

TIRF illumination intensity, *ί*, is described in terms of distance from the coverslip, *h,* by

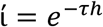

For simplicity, we measured the distance *h* in “layers”, with the depth of each layer corresponding to physical size of Akt-PH, which was estimated to be approximately 10 nm based on the sum of longest dimensions of Akt-PH and GFP in their respective crystal structures (PDB ID: 1UNQ and 1GFL). We solved for TIRF illumination intensity using the following values for our system: refractive indexes of solution *n_1_*=1.33 and coverslip *n_3_*=1.53, critical incidence angle θ_c_ =60.8 degrees. The laser wavelength used in our experiments was *λ_0_*=447 nm, and the experimental angle of incidence was θ_exp_=63 degrees. This produces a characteristic depth of *d_63_* =127 nm and an illumination decay of *τ_63_* =0.008 nm^-1^. We plot TIRF illumination intensity over distance in molecular layers and nanometers in Figure 2–figure supplement 2.

The values determined above allow us to estimate the contributions to our TIRF signal from the membrane vs. the cytosol. According to our calculation, the TIRF illumination intensity approaches 0 at around 500 nm, or layer *h_49_*. We consider the membrane and associated proteins to reside in layer *h_0_.* Under these conditions, at rest, 5% of total recorded TIRF fluorescence arises from *h_0_*, with the remainder originating from *h_1_-h_49_*. At rest, we assume that Akt-PH molecules are distributed evenly throughout layers *h_0_-h_49_*, with no Akt-PH bound to the membrane because the concentration of PIP_3_ in the PM is negligible at rest. Total fluorescence intensity measured before NGF application, *F_initial_,* depends on *m,* the number of molecules per layer at rest, B, the brightness of a single molecule of CFP, and TIRF illumination intensity, *ί*:

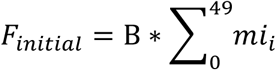

Normalizing our time traces to *F_initial_*, sets *F_initial_* = 1. We solved for *m* numerically using Excel (Microsoft, Redmond, WA; see Supplemental Excel File 1), and determined a value of 0.08. We assumed a fixed number of molecules in the field and that the only NGF-induced change was a redistribution of molecules among layers. The total fluorescence intensity measured after NGF application, *F_NGF_,* will reflect the redistribution of *Δm* molecules between membrane layer *h_0_* and all layers *h_0_-h_49_*, with free Akt-PH homogeneously distributed among these layers. Therefore, *F_NGF_* is a sum of fluorescence intensities of the number of bound molecules in the membrane layer *h_0_* and the free molecules in layers *h_1_-h_49_*:

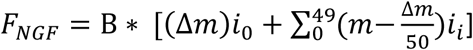

We solved for *Δm* using Excel, constraining F_NGF_ to the values we measured for control and TRPV1-expressing cells (data listed in the table in Figure 2–figure supplement 2B). Finally, we estimated the NGF-induced change in Akt-PH bound to the membrane as *R_m_*, the ratio of molecules in *h_0_* after NGF to that before NGF:

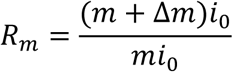

We compared R_m_ values to the F_NGF_ values listed in the table Figure 2–figure supplement 2B. For example, in cells expressing TRPV1, F_NGF_ of 1.54 led to 10 times more membrane-associated Akt-PH molecules. Note, that if we instead assume that the number of homogenously distributed molecules of Akt-PH does not change with NGF treatment, we calculate an Rm value of 8, very similar to the value of 10 obtained with redistribution of a fixed number of molecules. Both of these scenarios are independent of the initial Akt-PH fluorescence intensity in a given cell.

### Cell culture /transfection/ DNA constructs

F-11 cells (a gift from M.C. Fishman, Massachusetts General Hospital, Boston, MA; (Francel et al., 1987)) were cultured at 37°C, 5% CO2 in Ham’s F-12 Nutrient Mixture (#11765-054; Gibco) supplemented with 20% fetal bovine serum (#26140-079; Gibco, Grand Island, NY), HAT supplement (100 µM sodium hypoxanthine, 400 nM aminopterin, 16 µM thymidine; #21060-017; Gibco), and penicillin/streptomycin (#17-602E, Lonza, Switzerland). F-11 cells for imaging experiments were plated on Poly-Lysine (#P1274, Sigma, St. Louis, MO) coated 0.15mm × 25mm coverslips (#64-0715 (CS-25R15), Warner Instruments, Hamden, CT) in a 6-well plate. Cells were transfected with Lipofectamine 2000 (4ul/well, Invitrogen, Grand Island, NY) reagent using 1-3ug of cDNA per well. 24hrs post-transfection, media was replaced with HEPES-buffered saline (HBR, double deionized water and in mM: 140 NaCl, 4 KCl, 1 MgCl_2_, 1.8 CaCl_2_, 10 HEPES (free acid) and 5 glucose) for at least 2hrs prior to the imaging. During experiments, cells were treated with 100ng/ml NGF 2.5S (#13257-019, Sigma) or vehicle (HBR).

TRPV1-cYFP (rat) (Ufret-Vincenty et al., 2015), TRPV1-ARD-c tagRFP (rat), TRPV2-cYFP (rat) (Mercado et al., 2010) DNA constructs were made in the pcDNA3 vector (Invitrogen), where *“*-n” or “-c” indicates that the fluorescent protein is on the N- or C-terminus, respectively. TRPV4-EGFP (human) in pEGFP was obtained from Tim Plant (Charite-Universitatsmedizine, Berlin) (Strotmann et al., 2003). pEGFP-TRPM8 (rat) in pEGFP was obtained from Addgene (#64879, Addgene, Cambridge, MA). TRPM4-GFP (mouse) in the pEGFPN1 vector was obtained from Marc Simard (University of Maryland, College Park) (Gerzanich et al., 2009). TRPA1-GFP (zebrafish) in pcDNA5-FRT was obtained from Ajay Dhaka (University of Washington, Seattle) TrkA (rat) in the pcCMV5 vector and p75_NTR_ (rat) in the pcDNA3 vector were obtained from Mark Bothwell (University of Washington, Seattle). PH-Akt-cCerulean in the pcDNA3-k vector was made based on the construct in the pHR vector from the Weiner Lab (Toettcher et al., 2011). The function of the ion channels TRPV1, TRPV2, TRPV4, TRPM4, TRPM8, and TRPA1 was confirmed using Ca^2+^ imaging and/or patch clamp electrophysiology (data not shown).

### Western Blotting

For detection of relative expression of PI3K p85 alpha subunit, cells were transfected as described above for imaging experiments. 24hrs after transfection, cells were scraped off the bottom of 10 cm plates, washed with PBS 4 times and homogenized in Lysis buffer (1% Triton 25 mM Tris-HCl, 150mM NaCl, 1mM EDTA, pH7.4) for 2hrs with mixing at 4C. Lysates were spun down at 14000 rpm for 30min at 4C to remove the cell nuclei and debris. Cleared lysates were mixed with Laemmli 2x SDS sample buffer (#161-0737, Bio-Rad, Hercules, CA), boiled for 10 minutes and subjected to SDS PAGE to separate proteins by size. Gels were then transferred onto the PDVF membrane using Trans-Blot SD semi-dry transfer cell (Bio-Rad) at 15V for 50min. Membranes were blocked in 5% BSA TBS-T for 1hr and probed with primary antibody for 1hr at RT. Next, membranes were washed 6x times with TBS-T and probed with secondary antibodies conjugated with HRP for 1hr. After another set of 6 washes membranes were developed by addition of the SuperSignal^TM^ West Femto HRP substrate (#34096, Thermo, Grand Island, NY) and imaged using CCD camera-enabled imager. For quantification, blot images were analyzed in ImageJ. ROIs of the same size were drawn around the bands for p85 and tubulin, then mean pixel intensity was measured. Mean p85 intensities were normalized by dividing by mean tubulin intensities and plotted in Figure 3–figure supplement 1. Experiments were repeated with n=5 independent samples. Primary antibodies used were: anti-PI3K (alpha) polyclonal (#06-497 (newer Cat#ABS234), Upstate/Millipore, Burlington, MA) at 1:600 dilution; β Tubulin (G-8) (#sc-55529, Santa Cruz, Dallas, TX) at 1:200 dilution. Secondary antibodies used: Anti-Rabbit IgG (#074-1506, KPL/SeraCare Life Sciences, Milford, MA) at 1:30,000 dilution; Anti-Mouse IgG (#NA931, Amersham/ GE Healthcare Life Sciences, United Kingdom) at 1:30,000 dilution.

For detection of phosphorylated Akt, cells plated in 6-well plates were treated for the indicated amount of time (Figure 4, Figure 4–figure supplement 1) in the CO_2_ incubator at 37°C. Immediately after treatment, wells were aspirated and scraped in ice-cold lysis buffer (H_2_O, TBS,1% NP-40, 5mM NaF, 5mM Na_3_VO_4_, Protease inhibitors (#P8340, Sigma), Phosphatase Inhibitor Cocktail 2 (#P5726, Sigma). After incubation on ice for 15 min, lysates were cleared by centrifugation at 15k g for 15 min at 4°C. Protein contents of cleared lysates were measured using the BCA assay (#23225 Pierce) according to manufacturer’s protocol. Volumes of lysates were adjusted according to these measurements and subjected to SDS-PAGE. Gels were transferred onto PVDF membranes using wet-transfer. Membranes were blocked in TBS-T with 5% milk for 1hr and incubated overnight at 4°C with one of the following primary antibodies: pAKTs473 clone D9E (#4060, Cell Signaling), pAKTt308 clone 244F9 (#4056, Cell Signaling) or panAKT clone 40D4 (#2920, Cell Signaling) at 1:2500 dilution. Further procedures were as indicated in the previous paragraph. Data was normalized by diving the average intensity of a band by the average intensity of a blot and then dividing by that of a pan-Akt blot (Figure 4).

## Results

### NGF induces production of PIP_3_ by PI3K followed by trafficking of TRPV1 channels to the PM

It has been previously established that PI3K activity is required for NGF-induced trafficking of TRPV1 to the PM (Bonnington and McNaughton, 2003, Stein et al., 2006), but the role of PIP_3_, the product of PI3K activity, was unclear. We hypothesized that PIP_3_ constitutes part of the required signal for TRPV1 trafficking (Fig 1), as it does for trafficking of membrane/membrane proteins in other systems (Cheatham et al., 1994, Martin et al., 1996, Xu et al., 2016). To test this hypothesis, we used two-color TIRF microscopy to examine PIP_3_ production and TRPV1 trafficking simultaneously. We used F-11 cells transiently transfected with TRPV1 (rat) fused to YFP (referred to as TRPV1) and the NGF receptor subunits TrkA and p75_NTR_ (referred to as TrkA/ p75_NTR)._ In addition, the cells were co-transfected with a fluorescent probe that recognizes PIP_3_, the PIP_3_-specific Pleckstrin homology (PH) domain (James et al., 1996) from the enzyme Akt fused to CFP (referred to as Akt-PH (Frech et al., 1997)). TIRF microscopy isolates ^~^100 nm of the cell proximal to the coverslip (Ambrose,1961, Axelrod,1981), capturing the PM-proximal fluorescent signals. A change in Akt-PH fluorescence reflects a change in PIP_3_ concentration at the PM, thus serving as an indirect measure of PI3K activity. A change in TRPV1 fluorescence reflects a change in the number of TRPV1 channels at the PM.

To study NGF-dependent changes in PM Akt-PH and TRPV1, we alternated image acquisition of the Akt-PH and TRPV1 fluorescence signals (Figure 2A-D) before, during, and following a 10-minute exposure of the cells to NGF (100ng/ml) via its addition to the bath (bar with gray shading in Figure 2B, C). TIRF images of the same cell are shown for both TRPV1 and Akt-PH fluorescence signals at time points before (time point 1) and during (time point 2) NGF treatment. Figure 2A, top panel shows representative TIRF images of Akt-PH fluorescence of the individual F-11 cell footprint and Fig. 2A, bottom shows the corresponding signal for TRPV1. At rest, there was little TIRF fluorescence signal from Akt-PH domain from the footprint of the cell, which reflects to the low resting PIP_3_ levels at the PM (Haugh et al., 2000) (fig. 2A, 1 top panel). Upon addition of NGF, both PIP_3_ and TRPV1 levels at the PM increased, with time point 2 depicting the cell footprint intensity at steady state. For every cell, we normalized the mean fluorescence intensity within the footprint at each time point to the mean between 0-60 seconds prior to the application of NGF. The signals for Akt-PH and TRPV1 for the cell in 2A are shown in Figure 2B, and the collected data, showing the mean and standard error of the mean, are illustrated in Figure 2C.

**Figure 2.**
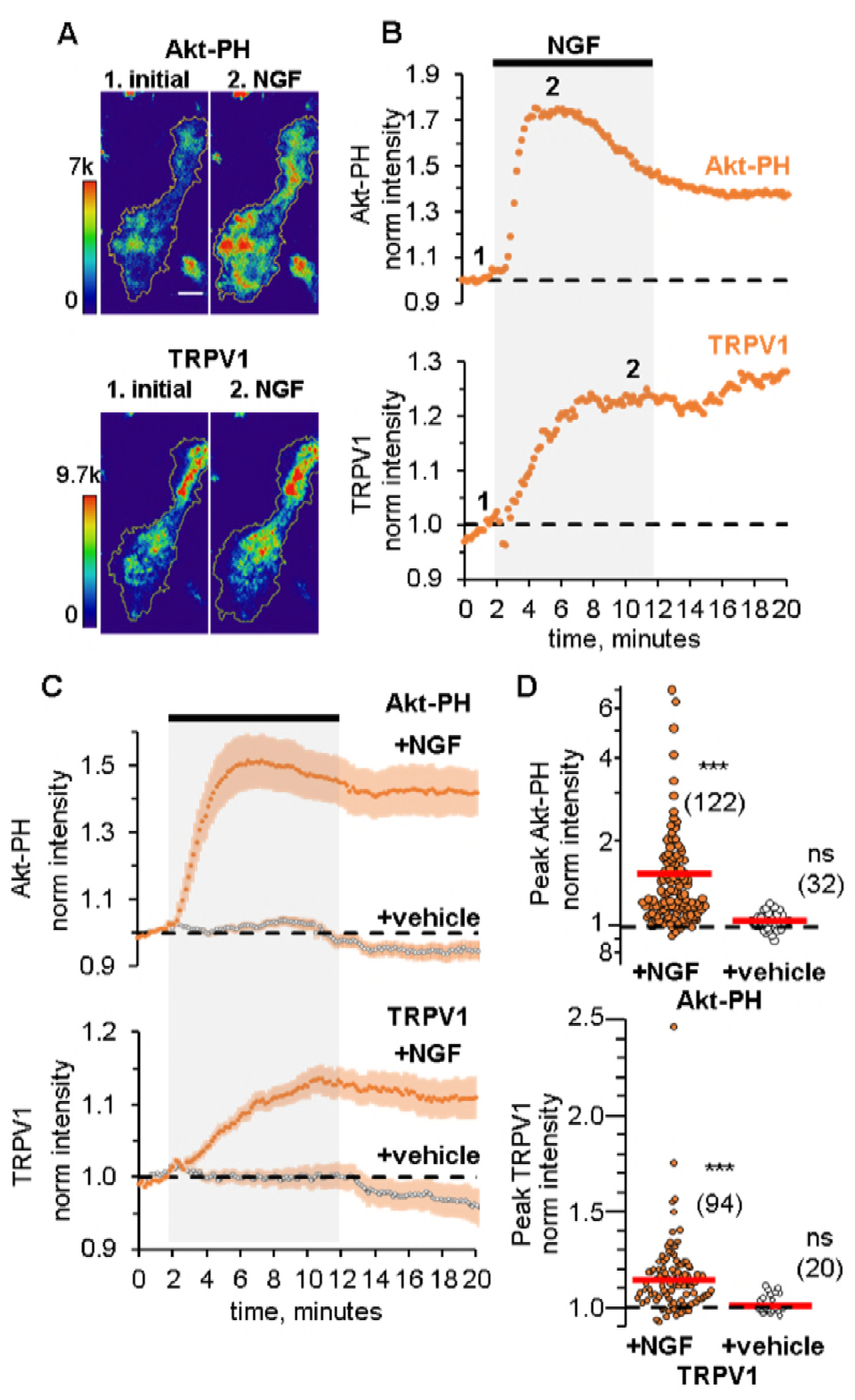
NGF increases PIP_3_ and recruits TRPV1 to the PM. (A) TIRF images of a representative F-11 cell transfected with TrkA/p75_NTR_, TRPV1 and Akt-PH. Images labeled 1 were collected before NGF application and those labeled 2 were collected at the plateau during NGF application, as indicated by the time points labeled in B. Scale bar is 10 µm. LUT bars represent background-subtracted pixel intensities. The yellow border represents the outline of the cell footprint. (Top) Fluorescence intensity from Akt-PH. (Bottom) Fluorescence intensity from TRPV1. (B) Time course of NGF-induced changes in fluorescence intensity for the cell shown in A. NGF (100 ng/mL) was applied during the times indicated by the black bar/gray shading. Intensity at each time point was measured as the mean gray value within the footprint (yellow outline in A). Data were normalized to the mean intensity values during the two minutes prior to NGF application. (C) And (D) Collected data for the group of cells tested. (C) Time course of NGF-induced changes in fluorescence intensity. Averaged time courses of TIRF intensity normalized as in B. Cells treated with either NGF (filled symbols) or vehicle (open symbols), as indicated. TRPV1 (bottom, n=94) and Akt-PH (top, n=122). Error bars are SEM (D) NGF-induced change in fluorescence intensity. Cells treated with NGF are represented by filled symbols and cells treated with vehicle are shown in open symbols. Averaged normalized TIRF intensity during NGF application (6-8 minutes for Akt-PH and 10-12 minutes for TRPV1). The red bars indicate the mean (see Table 1 and Table 2 for values). The y-axis is logarithmic. Asterisks indicate significance (paired Student’s t-test, p <0.001, see Table 1 and 2 for values), comparing each cell’s normalized intensity before and during NGF treatment.

Treatment of cells with NGF produced an increase in plasma-membrane associated Akt-PH, indicating that PIP_3_ levels in the PM increased. The increase was relatively rapid (approximately 4 min to peak) and partially decreased over time even in the continued presence of NGF (Figure 2 B, C, top), possibly due to TrkA/p75_NTR_ receptor internalization (Grimes et al., 1996, Ehlers et al., 1995). In contrast, NGF treatment increased the PM TRPV1 signal more slowly (approximately 8 min to peak) without apparent reversal to baseline over the duration of our experiments (Figure 2 B, C, bottom). The peak levels of Akt-PH and TRPV1 for all cells, represented as the normalized intensities measured at 4-6 minutes (for Akt-PH) and 8-10 minutes (for TRPV1) after the start of NGF application, are shown in the scatterplot of Figure 2D. The distributions were not normal, but skewed towards larger values. This distribution shape is characteristic of NGF-induced TRPV1 sensitization reported previously in DRG neurons, indicating that our cell expression model behaves similarly to isolated DRG neurons (Stein et al., 2006, Bonnington and McNaughton, 2003). NGF induced a significant increase in Akt-PH levels (Mean ± SEM: 1.54 ± 0.08 (n=122) compared to 1.01 ± 0.01 (n=32), Wilcoxon rank test p<0.001) compared to vehicle (Fig 2C, top panel, filled and open symbols respectively, paired t-test within each condition, see also Figure 2–figure supplement 1), and a significant increase in TRPV1 levels compared to vehicle (Mean ± SEM: 1.15 ± 0.02 (n=94) compared to 0.99 ± 0.01 (n=20), Wilcoxon rank test p<0.001; Fig2 C, bottom panel, filled and open symbols respectively, see also Figure 2–figure supplement 1).

TIRF microscopy is often discussed as a method that isolates a fluorescence signal at the PM (Axelrod, 1981). Indeed, illumination falls off exponentially with distance from the coverslip (Ambrose, 1961). Nevertheless, with a typical TIRF setup such as that used for this study (see Materials and Methods) ̃90% of the signal comes from the cytosol (Figure 2–figure supplement 2, also see Methods), assuming the incidence of light was at the critical angle and that the membrane bilayer and associated protein layer extends up to ̃10 nm from the coverslip. The contamination of the TIRF signal with fluorescence from the cytosol leads to an underestimation of the change in PM-associated fluorescence from Akt-PH and TRPV1. Under our experimental conditions, we estimate that the ratio of the total fluorescence intensity measured after and before NGF application, F_NGF_, of 1.54 translates into ̃10-fold increase in PM-associated fluorescence, R_m_ (Figure 2–figure supplement 2; see Methods).

### TRPV1 potentiates NGF-induced PI3K activity

We compared NGF-induced PI3K activity in control cells to that in TRPV1-transfected cells. Remarkably, we found that, TRPV1 appeared to increase the activity of PI3K. Figure 3A shows the average normalized Akt-PH fluorescence intensity over time from control cells. In non-TRPV1 expressing cells, NGF caused an increase in the normalized Akt-PH fluorescence of 1.08 ± 0.03 (n=75). Figure 3B shows Akt-PH fluorescence in cells that were co-transfected with TRPV1 (same data as in Fig2C). Co-expression of TRPV1 caused NGF-induced Akt-PH translocation to increase to 1.54 ± 0.08 (n=122). As seen from the scatter plot in Figure 3C, TRPV1-expressing cells produced higher levels of PIP_3_ compared to control cells. A Student’s t-test assuming unequal variances showed that the difference between control and TRPV1-expressing populations was statistically significant at an alpha of 0.001 (see Table 1 for values). It should be noted that the dynamics of PIP_3_-generation was also different: in the absence of TRPV1, NGF-induced increases in PIP_3_ levels were sustained, whereas in TRPV1-expressing cells there was an initial rise followed by a slight decrease in PIP_3_ levels.

**Figure 3.**
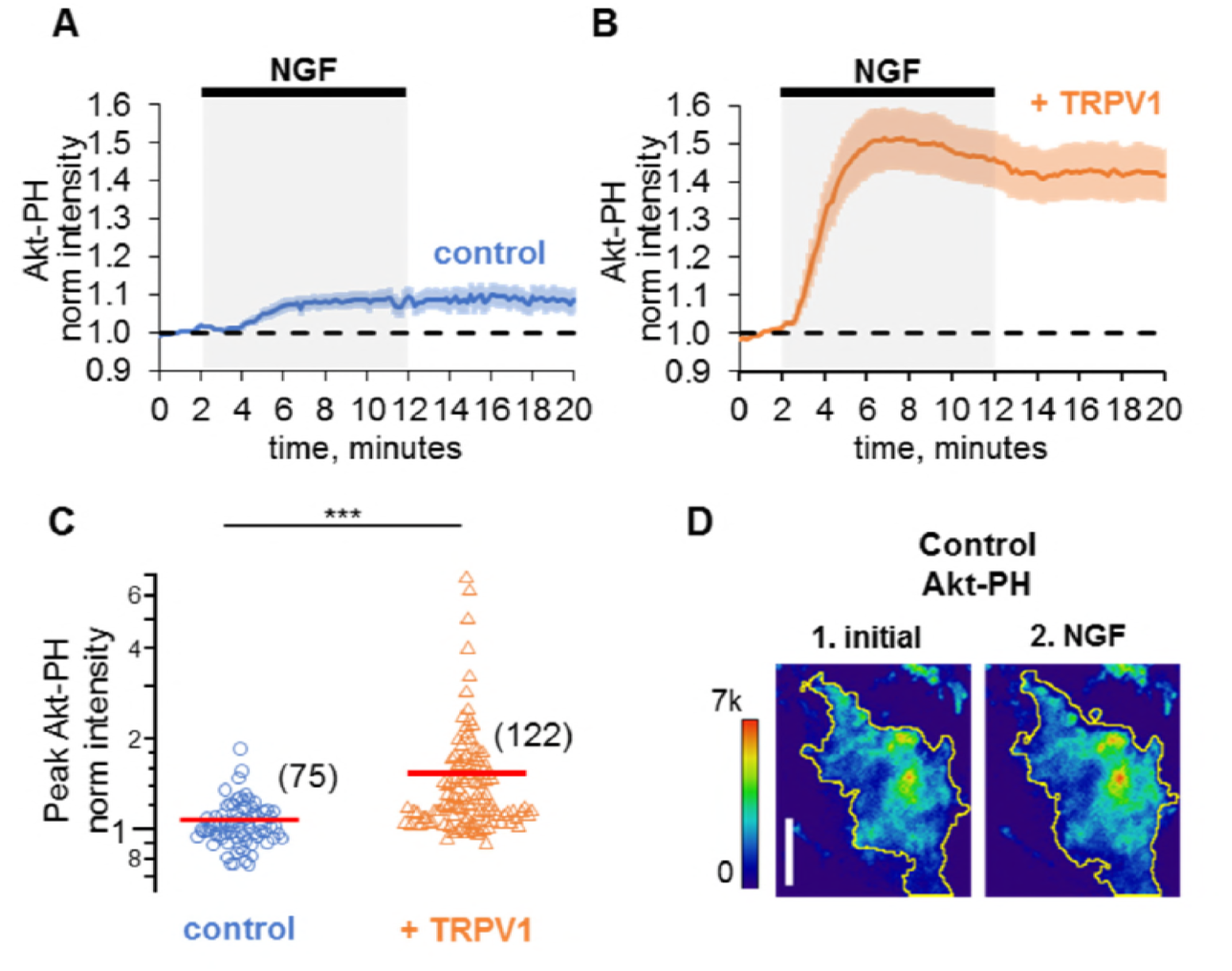
TRPV1 potentiates NGF-induced PIP_3_ production by PI3K. (A) And (B) Collected data for the NGF-induced change in Akt-PH. Traces represent the Akt-PH mean normalized fluorescence intensity and error bars represent the SEM. (A) NGF induces moderate activation of PI3K in control cells transfected with TrkA/p75_NTR_ and Akt-PH, but not TRPV1; n=75. (B) NGF induces high PI3K activity in cells expressing TRPV1 in addition to TrkA/p75_NTR_ and Akt-PH; n=122. Same data as in Fig. 2C (C) NGF-induced changes in Akt-PH fluorescence intensity for control cells (circles) and cells expressing TRPV1 (triangles). Averaged normalized TIRF intensity during NGF application (6-8 minutes). Red bars indicate mean (see Table 1 for values). The y-axis is logarithmic. Asterisks indicate significance of Student’s t-test assuming unequal variances, one-tail p value <0.001. (D) TIRF images of a representative control F-11 cell transfected with TrkA/p75_NTR_ and Akt-PH, no TRPV1. Images labeled 1 were collected before NGF application and those labeled 2 were collected at the plateau during NGF application, as indicated by the time points labeled in Fig2B. Scale bar is 10 µm. LUT bars represent background-subtracted pixel intensities. The yellow border represents the outline of the cell footprint.

**Table 1.** Normalized Akt-PH fluorescence intensities measured during NGF application for all discussed conditions. The number of cells in the data set collected over at least three different experiments is given by n. For all columns, asterisks indicate significance at alpha of 0.05.

| Akt-PH from | NGF<br>Mean ±<br>SEM | n= | Paired t-test,<br>p value <sup>1</sup> | t-test<br>compared to<br>control,<br>p value <sup>2</sup> | t-test<br>compared to<br>TRPV1,<br>p value <sup>3</sup> |
| --- | --- | --- | --- | --- | --- |
| control | 1.08 ± 0.03 | 75 | 0.0001 * | - | - |
| TRPV1 | 1.54 ± 0.8 | 122 | 10<sup>-10</sup> * | 10<sup>-7</sup> * | - |
| TRPV1-ARD | 1.32 ± 0.2 | 80 | 10<sup>-8</sup> * | 10<sup>-5</sup> * | 0.01 |
| TRPV2 | 1.23 ± 0.18 | 61 | 10<sup>-5</sup> * | 0.008 * | 0.001 * |
| TRPV4 | 1.28 ± 0.14 | 29 | 0.0002 * | 0.006 * | 0.008 * |
| TRPM4 | 1.20 ± 0.09 | 63 | 10<sup>-7</sup> * | 0.003 * | 0.0001 * |
| TRPM8 | 1.07 ± 0.02 | 73 | 0.0002 * | 0.29 | 10<sup>-8</sup> * |
| TRPA1 | 1.14 ± 0.05 | 59 | 10<sup>-6</sup> * | 0.05 | 10<sup>-6</sup> * |

<sup>1</sup> Student's paired t-test was performed on data for each condition individually before and after NGF.
<sup>2</sup> Student's t-test assuming unequal variances for each condition and control, one-tail p value (alpha of 0.05 for 7 comparisons was post-hoc adjusted to 0.007).
<sup>3</sup> Student's t-test assuming unequal variances for each condition and TRPV1, one-tail p value (alpha of 0.05 for 6 comparisons was post-hoc adjusted to 0.008),

One possible cause for the potentiation of NGF-induced PI3K activity we observed in TRPV1- expressing cells could be a change in PI3K expression levels in TRPV1 vs. control cells. To determine whether this was the case, we performed Western blot analysis with an anti-p85α antibody to quantify the PI3K protein levels across transfection conditions. As shown in Figure 3–figure supplement 1, expression of TRPV1 did not alter the expression level of the p85α subunit of PI3K. We quantified protein expression levels using densitometry, and normalized expression to tubulin, giving the relative expression levels shown in Figure 3–figure supplement 1. Average relative p85α expression levels were similar between non-TRPV1 expressing cells and cells expressing TRPV1 (n=5, Student’s t-test p value was 0.95). We conclude that a difference in PI3K expression in TRPV1-expressing vs. control cells did not account for the observed TRPV1-induced potentiation of NGF-stimulated PI3K activity.

TRPV1 potentiation of NGF-induced PI3K activity should enhance other PIP_3_-dependent downstream signaling processes. Phosphorylation of the protein kinase Akt (also known as PKB) is a well-studied signaling event downstream of PIP_3_ production (Burgering and Coffer, 1995, Kohn et al., 1995). Akt is phosphorylated in a PIP_3_-dependent manner at two sites, T308 and S473, by PDK1 (Alessi et al., 1997, Stokoe et al., 1997) and mTORC2 respectively (Sarbassov et al., 2005). Phosphorylation of Akt at these two sites leads to full activation of Akt, regulating a variety of cellular processes. We therefore tested whether TRPV1 potentiates NGF-induced phosphorylation of Akt.

To determine whether Akt phosphorylation was enhanced in TRPV1-expressing cells, we performed Western Blot analysis using phospho-specific Akt antibodies, reprobing the blots with a pan Akt antibody for normalization purposes (Figure 4A). Because phosphorylation at T308 and S473 are differentially regulated, we used three concentrations of NGF (5, 25, and 100 ng/mL) and two incubation times (1 and 5 minutes). We observed increased phosphorylation at both T308 and S473 in TRPV1- expressing cells compared to control cells for almost all trials with all three NGF concentrations and both time points (Figure 4B, C). The enhanced NGF-induced Akt phosphorylation was statistically significant for both T308 and S473 sites (Figure 4D, E; paired Student’s t-test for T308 p = 0.02 and S473 p=0.008) for all conditions pooled together. Thus, TRPV1 potentiation of NGF-induced PI3K activity is sufficient to enhance PIP_3_ levels to increase Akt phosphorylation.

As a control, we examined whether NGF-induced phosphorylation at both T308 and S473 required expression of TrkA/p75_NTR_. NGF-induced phosphorylation of Akt was not observed in cells in which TrkA/p75_NTR_ were not expressed (Figure 4–figure supplement 1). Importantly, we confirmed that co-expression of Akt-PH, corresponding to the PIP_3_-binding domain only, did not alter NGF-induced Akt phosphorylation (Figure 4–figure supplement 1). Finally, Figure 4 shows that the extent of Akt phosphorylation in unstimulated cells was indistinguishable in control vs. TRPV1-expressing cells at both S308 (Intensity_pAkt/panAkt_: 0.075 ± 0.004 for control and 0.076 ± 0.004 for TRPV1, Mean ± SEM, n=3, paired Student’s t-test p = 0.95) and T473 sites (Intensity_pAkt/pan Akt_: 0.3 ± 0.24 for control and 0.23 ± 0.14 for TRPV1, Mean ± SEM, n=3, paired Student’s t-test p = 0.44), indicating that TRPV1 did not perturb the levels of PIP_3_ at rest. Together these data demonstrate that TRPV1 potentiation of NGF-induced PI3K activity can be observed at the level of downstream Akt phosphorylation.

**Figure 4.**
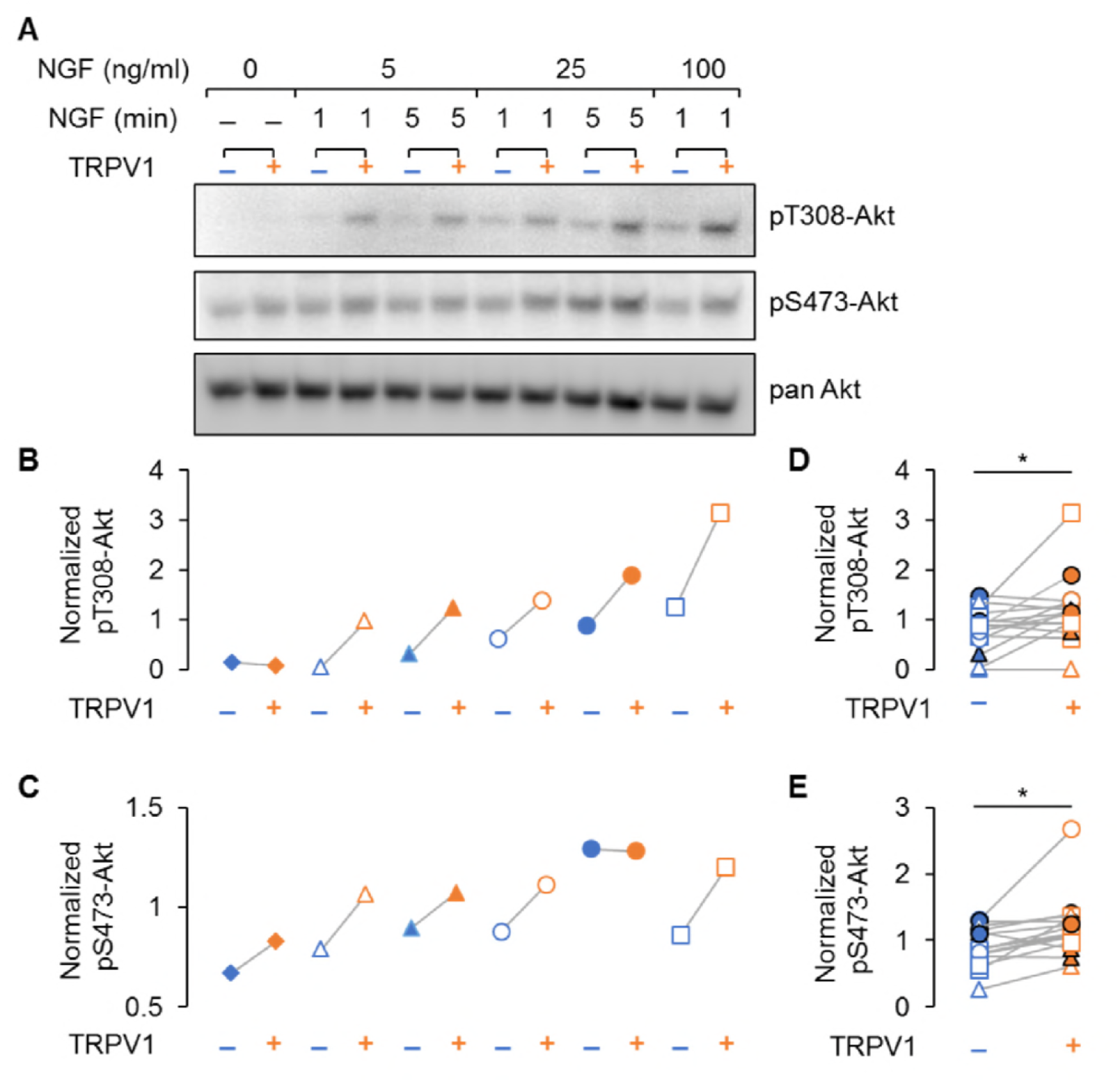
TRPV1 enhances NGF-induced Akt phosphorylation. (A) Representative immunoblot staining for analysis of Akt phosphorylation in F11 cells transfected same as in imaging experiments. Cells were treated with indicated dose of NGF for an indicated amounts of time, lysed and loaded on SDS-PAGE. The same membrane was probed with pAKTs473, stripped and re-probed with pAKTt308 and again with panAKT antibodies (See Methods). (B) and (C) Analysis of the representative blots shown in (A). Each band average intensity was normalized to the average of the blot and then divided by that of the corresponding lane of the pan- Akt blot. Akt phosphorylated at T308 (B) and S473 (C) from control cells (blue symbols) and cells expressing TRPV1(orange symbols) treated with NGF (5, 25 or 100ng/ml) for 1 or 5 minutes as indicated in (A). Triangles represent treatment with NGF 5 ng/ml, circles – 25 ng/m, squares – 100 ng/ml. Open symbols represent treatments for 1 min and filled symbols – 5 min. (D) and (E) Normalized phospho-Akt intensities from all indicated conditions are pooled together for the n=3 of independent experiments. Paired Student’s t-test for pT308-Akt p = 0.02 and for pS473-Akt p = 0.008.

Based on our data and the vast literature on PI3K, we can consider three levels of PI3K activity. Figure 4–figure supplement 2A shows a simplified cartoon representation of the signaling events at the PM in the absence of TRPV1. Resting cells have low PIP_3_ levels, because PI3K (green circle) is inactive in the absence of the NGF. Application of NGF (purple circle) causes auto-phosphorylation of TrkA/p75_NTR_ (brown oval), recruitment of PI3K to the PM and release of PI3K auto-inhibition, producing moderate PI3K activity levels. Figure 4–figure supplement 2B shows a cartoon representation of a cell co-expressing TRPV1 (orange). For these TRPV1-expressing cells, binding of NGF to TrkA/p75_NTR_ would activate PI3K and the TRPV1-interacting PI3K would have a higher level of activity than PI3K in cells not expressing TRPV1.

### Mechanism for potentiation of NGF-induced PI3K activity by TRPV1

Two possible mechanisms for NGF-induced potentiation of PI3K activity by TRPV1 are depicted in fig. 4A. In one mechanism, binding to the integral membrane protein TRPV1 may stabilize the association of PI3K with the membrane. The stabilization could occur at rest, in the absence of NGF, and/or upon NGF stimulation, after activated TrkA recruits PI3K to the membrane (Figure 5A, left). There are precedents for an association mechanism. A mutation in the p110 catalytic subunit of PI3K (H1047R) increases membrane association, thereby increasing activity (Gabelli et al., 2014, Carson et al., 2008). In addition, the membrane-bound version of the small GTPase H-Ras associates with PI3K, accelerating formation of a PI3K-phosphotyrosine complex and modestly slowing its off rate (Buckles et al., 2017, Burke and Williams, 2015). A second possible mechanism is that TRPV1 allosterically enhance the catalytic activity of NGF-activated PI3K (Figure 5A. right). The two mechanisms are not mutually exclusive, but do make different predictions about potentiation of PI3K by a soluble fragment of TRPV1 that is not bound to the PM. In the first case, we would expect that a TRPV1 fragment corresponding to the PI3K binding site would not be sufficient for potentiation, as it would not localize PI3K at the PM in resting cells. In the second case, the fragment would produce potentiation of PI3K activity via an allosteric mechanism.

**Figure 5.**
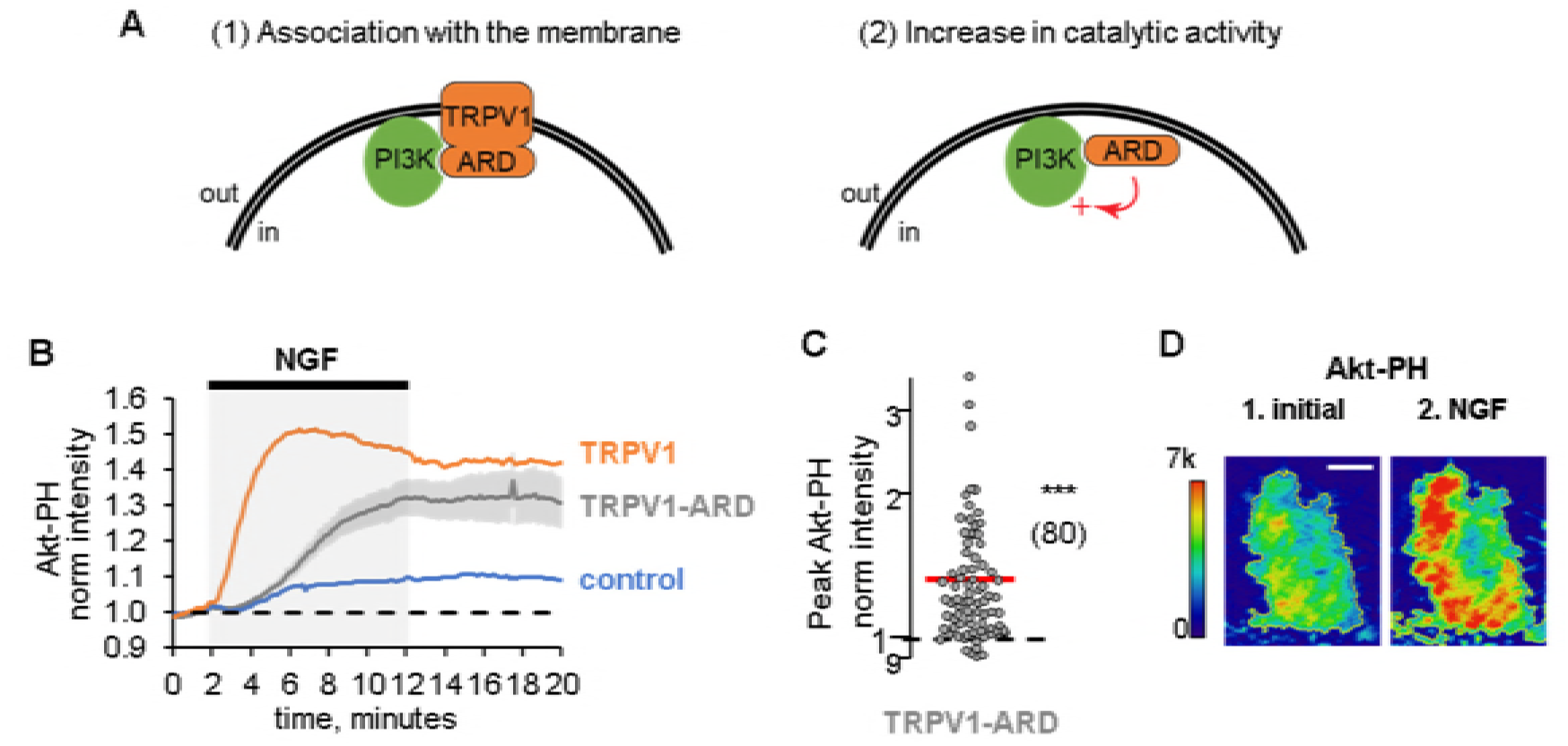
TRPV1-ARD potentiates NGF-induced PI3K activity. (A) Two possible mechanisms for PI3K potentiation by TRPV1-ARD. (1) TRPV1 may stabilize association of PI3K with the PM and/or (2) TRPV1-ARD may allosterically increase PI3K catalytic activity. (B) TRPV1-ARD potentiates NGF-induced PIP_3_ production by PI3K. Time course of NGF-induced changes in Akt-PH fluorescence intensity. NGF (100 ng/mL) was applied during the times indicated by the black bar/gray shading. Averaged normalized TIRF intensity from cells transfected with Akt-PH control (without TRPV1, n=75), TRPV1 (n=122), or TRPV1-ARD (n=80). Traces represent the mean and error bars represent the SEM. Control and TRPV1 data are the same as in Fig. 3A & B, error bars removed for clarity. (C) NGF-induced change in fluorescence intensity for Akt-PH (gray filled circles) in cells transfected with TRPV1-ARD. Averaged normalized TIRF intensity during NGF application (10-12 minutes). Red bars indicate mean values (see Table 1). The y-axis is logarithmic. Asterisks indicate significance (paired Student’s t-test, p <0.001), comparing each cell’s normalized intensity before and during NGF treatment. (D) TIRF images of a representative F-11 cell transfected with TrkA/p75_NTR_, Akt-PH and TRPV1-ARD. Images labeled 1 were collected before NGF application and those labeled 2 were collected at the plateau during NGF application, as indicated by the time points labeled in Fig2B. Scale bar is 10 µm. LUT bars represent background-subtracted pixel intensities. The yellow border represents the outline of the cell footprint.

We hypothesized that the Ankyrin repeats domain of TRPV1 (TRPV1-ARD) constitutes the PI3K binding site based the conserved role of ARDs as platforms for protein-protein interactions (Mosavi et al., 2004). ARDs are conserved and abundant protein domains that consist of multiple ̃33 amino acid helix-loop-helix Ankyrin repeats stacked together in a spring-like structure (Mosavi et al., 2004). The ARD of TRPV1, consisting of amino acids 111-359 (Figure 5–figure supplement 1) (Lishko et al., 2007), has been proposed to be involved in the modulation of channel function by interacting with intracellular regulators, including calmodulin, ATP, protein kinases and scaffolding proteins (Rosenbaum et al., 2004, Numazaki et al., 2003, Lishko et al., 2007, Morenilla-Palao et al., 2004, Zhang et al., 2008).

To determine whether the ARD is sufficient for potentiation of NGF-induced PI3K activity and whether potentiation requires localization at the PM, we expressed the ARD fragment (TRPV1-ARD) and measured NGF-induced PI3K activity using Akt-PH (Fig. 5B). Consistent with the allosteric model in Fig. 5A, potentiation of NGF-induced PI3K activity was observed in cells that expressed TRPV1-ARD (Fig. 5B, C; Mean ± SEM: 1.32 ± 0.02 (n=80); control vs TRPV1-ARD: Student’s t-test, p=10^−5^). The kinetics of this potentiation were somewhat slower with TRPV1-ARD compared to TRPV1, so that Akt-PH reached steady-state levels somewhat later during NGF treatment (Fig. 5B). Nevertheless, the ARD fragment fully reconstituted the amplitude of potentiation observed with full-length TRPV1 (TRPV1-ARD vs TRPV1: Student’s t-test, p=0.01), indicating that membrane localization was not required. Together, our data indicate that potentiation of NGF-induced PI3K activity by TRPV1 is mediated by its ARD and that the mechanism of potentiation likely involves an allosteric stimulation of the catalytic activity of PI3K by its direct interaction with the TRPV1 ARD. Whether the slower NGF-induced increase in Akt-PH observed with TRPV1-ARD compared with full-length TRPV1 (Fig. 3B) arises from a missing component of association of PI3K with the membrane remains to be determined.

### Potentiation of PI3K and NGF-induced trafficking are conserved among TRPV channels

The conservation of structure within the ARD of TRPV channels (Figure 5–figure supplement 1) raised the question of whether reciprocal regulation of PI3K activity and channel trafficking may be conserved among TRPV channels as well. To test this hypothesis, we examined NGF-induced PI3K activity in cells expressing other ARD-containing TRP channels. We tested two additional TRPV channels, TRPV2 (rat) and TRPV4 (human), whose ARD structures are very similar to that of TRPV1 (Figure 5–figure supplement 1). The NGF-induced increase in Akt-PH at the membrane was statistically significant compared to control cells for both TRPV2 and TRPV4 (Figure 6A, see also Table 1 for values). In addition, using TRPV2 and TRPV4 fused to fluorescent proteins, we found that they were both trafficked to the PM in response to NGF (Figure 6B, Figure 6–figure supplement 1, 2 and Table 1). We conclude that the ARD of TRPV1, TRPV2, and TRPV4 all support potentiation of NGF-induced PI3K activity and traffic to the PM in response to NGF.

**Figure 6.**
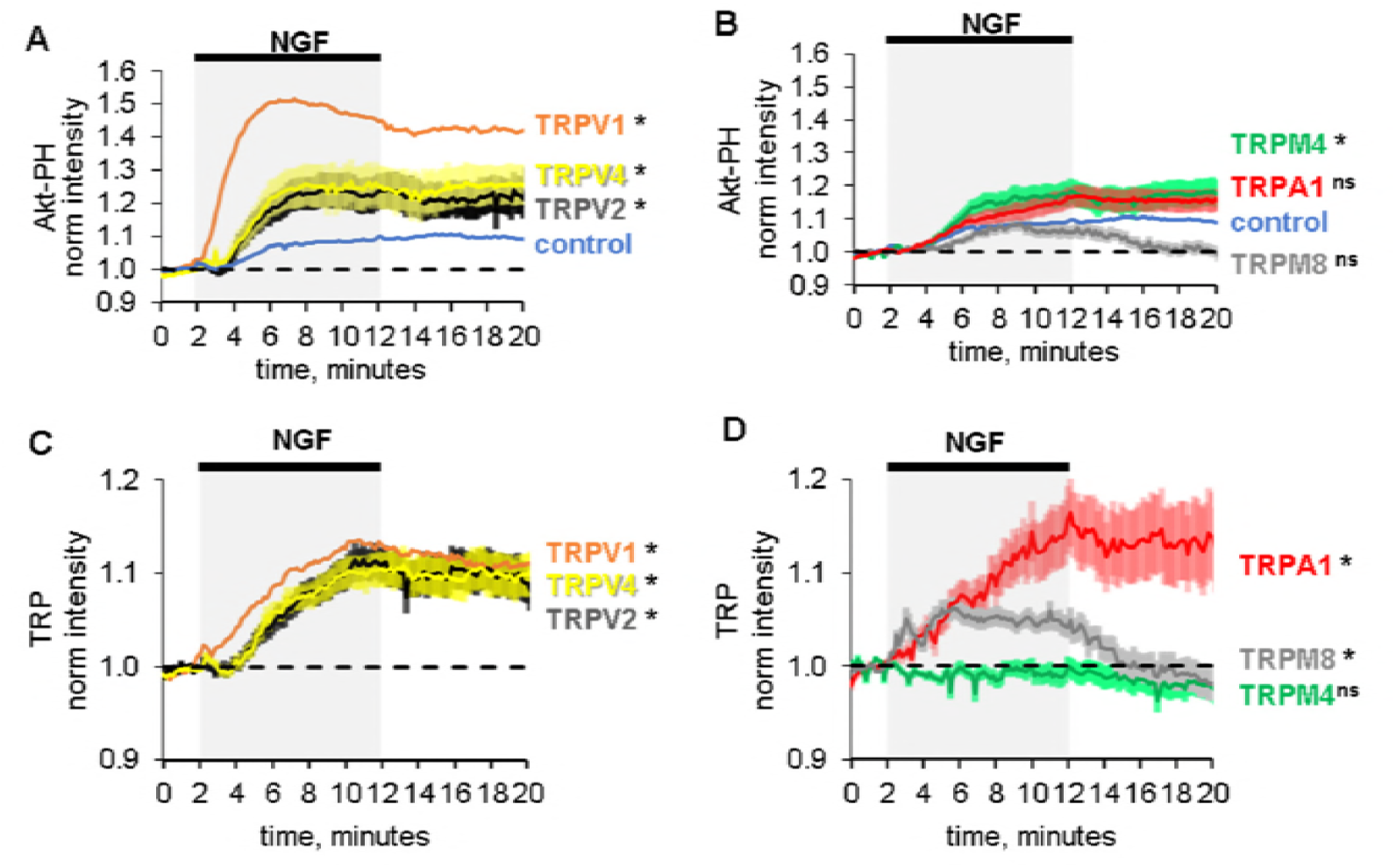
Potentiation of PI3K and NGF-induced trafficking are conserved among TRPV channels. Time course of NGF-induced changes in fluorescence intensity. NGF (100 ng/mL) was applied during the times indicated by the black bar/gray shading. Traces represent the mean, error bars are SEM. Control and TRPV1 data same as in Fig. 3A & B with error bars removed for clarity. (A-B) Averaged normalized TIRF intensity of Akt-PH from cells transfected with TrkA/p75_NTR_ and Akt-PH and: (A) no channel (control; blue; n=75); TRPV1 (orange; n=122); TRPV2 (black; n=61); TRPV4 (yellow; n=29) (B) no channel (control; blue; n=75); TRPM4 (n=63); TRPM8 (n=73); TRPA1 (n=59). (C-D) Averaged normalized TIRF intensity of individual TRP channels. Color scheme as in (A-B) with the cell numbers as follows: (C) TRPV1 (n=94); TRPV2 (n=62); TRPV4 (n=48); (D) TRPM4 (n=65); TRPM8 (n=67); TRPA1 (n=52).

We next addressed whether reciprocal regulation of PI3K activity and channel trafficking extends to TRPA1, which contains an N-terminal ARD with a very different structure compared to the TRPV channels (Figure 5–figure supplement 1). TRPA1 (zebrafish) appeared to potentiate NGF-induced PI3K activity (Figure 6B), although this potentiation did not reach statistical significance (Mean ± SEM: 1.14 ± 0.05, p value=0.05). Poor expression of TRPA1 may contribute to the apparent lack of significance in potentiation of NGF-induced PI3K activity (Mean ± SEM: 1518 ± 321 au for TRPA1 compared to 3039 ± 296 au for TRPV1). In addition, TRPA1 was trafficked to the PM in response to NGF (Figure 6D). We favor an interpretation in which the ARD of TRPA1 does somewhat potentiate NGF-induced PI3K activity at least to the extent required for its trafficking to the PM. More sensitive cell assays for PIP_3_ and in vitro studies will be required to test this hypothesis.

We also tested two other TRP channels, TRPM4 (mouse) and TRPM8 (rat), that do not contain ARDs. We found that they did not show reciprocal regulation with both PI3K activity and channel trafficking. Although TRPM4 channels potentiated NGF-induced PI3K activity (Figure 6C, Figure 6–figure supplement 1, 2, see also Table 1 for values), they did not traffic to the PM in response to NGF (Figure 6D, Figure 6–figure supplement 1, 2, see also Table 2 for values). TRPM8 channels did not potentiate PI3K activity, although they transiently trafficked to the PM in response to NGF. These data suggest that whereas the reciprocal regulation of PI3K activity and channel trafficking are conserved among TRPV channels with conserved ARD structures, this conservation does not extend to other TRP channels with different N-terminal domain structures.

**Table 2.**
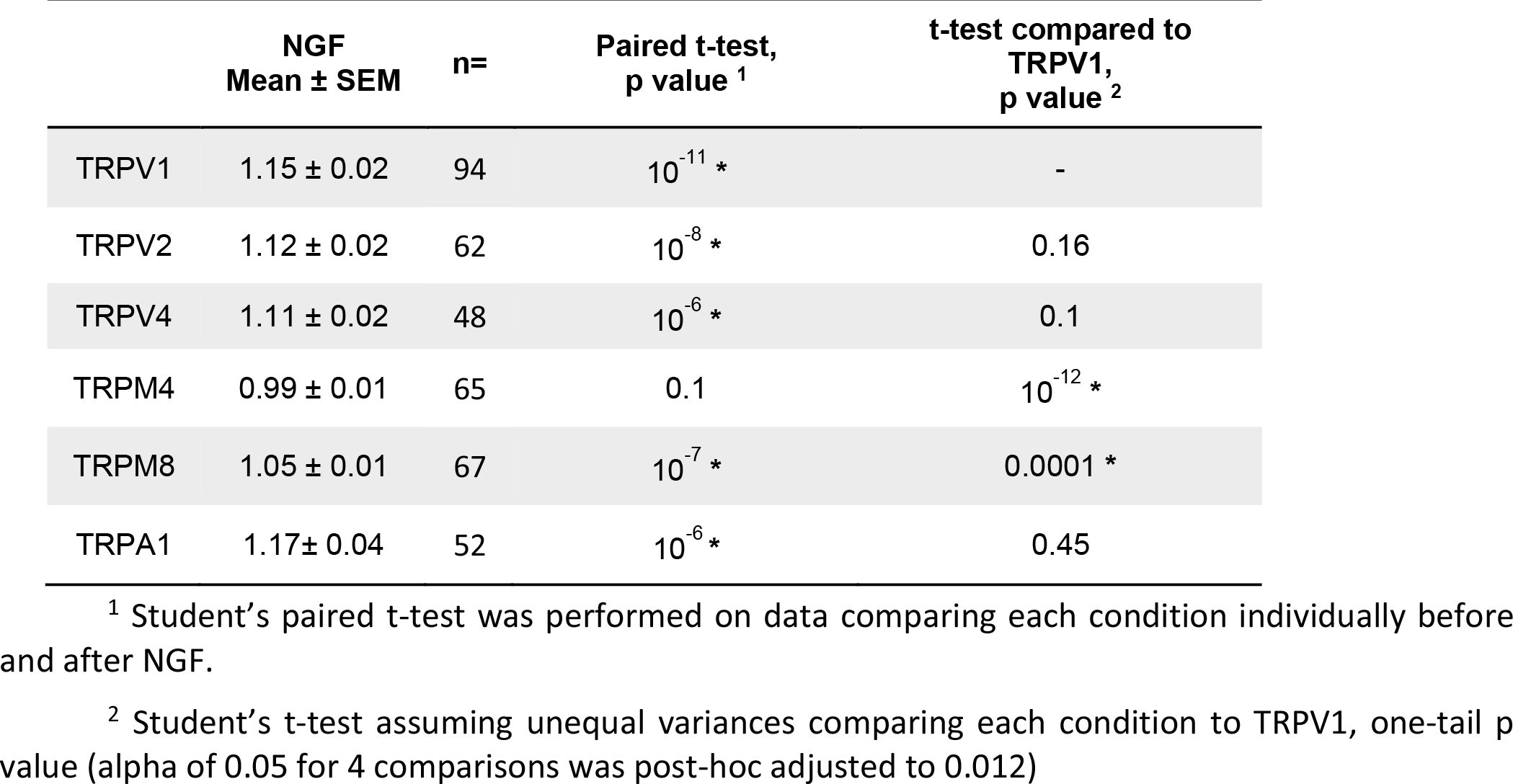
Normalized TRP channel fluorescence intensities measured during NGF application for all discussed conditions. The number of cells in the data set collected over at least three different experiments is given by n. For all columns, asterisks indicate significance at alpha of 0.05.

## Discussion

Using dual color TIRF imaging, we demonstrate that stimulation of TrkA/p75_NTR_ receptor with NGF rapidly increases levels of the signaling lipid PIP_3_. Increased PIP_3_ levels are followed by an increase in TRPV1 at the PM. Surprisingly, expression of TRPV1 potentiates NGF-induced production of PIP_3_ by PI3K. This potentiation occurs without any change in in PI3K expression. We previously showed that the N-terminus of TRPV1 interacts directly with PI3K. We localized the PI3K binding site to the ARD of TRPV1, and found that a soluble fragment corresponding to the ARD was sufficient for potentiation of NGF-induced PI3K activity. Other TRPV channels also potentiated NGF-induced PI3K activity and were trafficked to the PM in response to NGF. Our data reveal a new mechanism of reciprocal regulation among TRPV channels and PI3K that is mediated by allosteric regulation of PI3K by the TRPV ARDs.

Our conclusions are summarized in the illustration of Figure 7. In cells that do not express TRPV1, NGF-induced PI3K activity is moderate, generating moderate levels of PIP_3_ (Figure 7A). We speculate that these moderate levels of PIP_3_ are not sufficient to support increased trafficking of membrane protein-containing vesicles to the PM. In contrast, co-expression of TRPV1 potentiates NGF-induced PI3K activity, generating higher levels of PIP_3_ (Figure 7B). We propose that these higher levels of PIP_3_ are necessary to support trafficking of TRPV1-continaing vesicles to PM.

**Figure 7.**
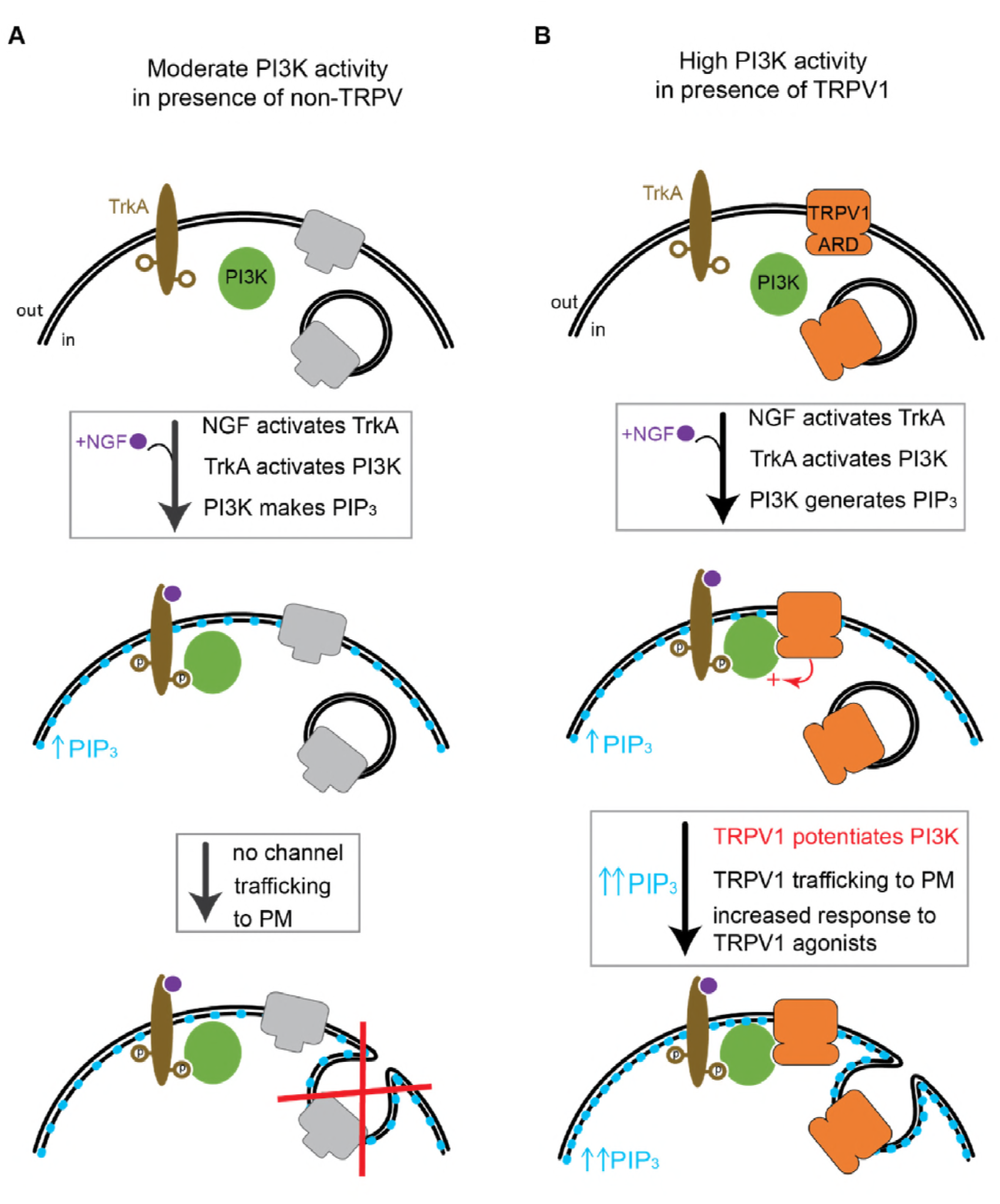
Cartoon representation of reciprocal regulation between TRPV channels and PI3K. (A) We speculate that moderate PI3K activity levels in presence of the non-TRPV channel are not sufficient to promote channel trafficking to the PM. (B) In presence of TRPV1 (and other TRPV channels) PI3K activity is potentiated which results in sufficient PIP_3_ levels for channel trafficking to the PM.

Increased trafficking of TRPV1 to the cell surface is essential for sensitization to noxious stimuli produced by NGF and other inflammatory mediators (Morenilla-Palao et al., 2004, Ferrandiz-Huertas et al., 2014). Although the involvement of PI3K in NGF-induced sensitization has been known for over a decade (Bonnington and McNaughton, 2003, Stein et al., 2006), the role, if any, of direct binding of TRPV1 and PI3K was unclear. Here we show that ARD region of TRPV1 that binds PI3K potentiates NGF-induced PI3K activity. Furthermore, our data suggest that without TRPV1 potentiation of PI3K, NGF signaling would not produce sufficient PIP_3_ to promote channel trafficking. This hypothesis is supported by our finding that, of the TRP channels tested here, only those that potentiate NGF-induced PI3K activity also traffic to the PM in response to NGF. Interestingly, TRPM4 seemed to be the exception, although it may be involved in other mechanism for PI3K potentiation, it did not exhibit channel trafficking. Future studies that decouple potentiation of PI3K activity from the expression of specific TRPV channels will be needed to determine whether the reciprocal regulation between ARD-containing TRPV channels and PI3K serves an obligate role in channel sensitization.

Is reciprocal regulation among TRPV channels and PI3K relevant beyond pain signaling? TRPV channels have been proposed to be involved in RTK/PI3K signaling in a variety of cell types (Reichhart et al., 2015, Katanosaka et al., 2014, Jie et al., 2015, Sharma et al., 2017). For example, TRPV2 is co-expressed in muscle cells with the insulin like growth factor receptor and is known to be important in muscle loss during muscular dystrophy (Iwata et al., 2003). The mechanism is believed to involve insulin like-growth factor receptor activation leading to increased trafficking of TRPV2 to the sarcolemma, Ca^2+^ overload/cytotoxicity, and cell death (Iwata et al., 2003, Perálvarez-Marín et al., 2013, Katanosaka et al., 2014). Whether TRPV2 potentiates insulin like growth factor-induced PI3K activity remains to be determined. The co-expression of TRPV channels with RTK/PI3K in other tissues, including nerve (TRPV1/NGF) (Tanaka et al., 2016), muscle (TRPV2/IGF) (Katanosaka et al., 2014) and lung (TRPV4/TGFβ1) (Rahaman et al., 2014) raises the question of whether reciprocal regulation among TRPV channels and PI3K plays a role in RTK signaling in cell development, motility, and/or pathology.

## Acknowledgements

We would like to thank Mika Munari, Eric Senning, Gilbert Martinez, Mark Bothwell, Bertil Hille, William N. Zagotta, Shao-En Ong, Tamara Rosenbaum and Gaby Bergollo for helpful discussions and Mario Rosasco for making Figure 5–figure supplement 1. We are grateful to the following individuals for providing cDNA constructs: Tim Plant (Charite-Universitatsmedizine, Berlin) for TRPV4; Marc Simard (University of Maryland, College Park) for TRPM4; Mark Bothwell (University of Washington, Seattle) for TrkA and p75_NTR_; Ajay Dhaka (University of Washington) for TRPA1; and Orion Weiner (UCSF) for PH-Akt.

Research reported in this publication was supported by the National Eye Institute of the National Institutes of Health under award numbers R01EY017564 (to SEG), by the National Institute of General Medical Sciences of the National Institutes of Health under award numbers R01GM100718 and R01GM125351 (to SEG), by the National Institute of Mental Health under award number R01MH113545 (to SEPS), by the National Institute of Biomedical Imaging and Bioengineering of the National Institutes of Health under award number T32EB001650 (to AS), and by the following additional awards from the National Institutes of Health: S10RR025429, P30DK017047, and P30EY001730. The content is solely the responsibility of the authors and does not necessarily represent the official views of the National Institutes of Health. The authors declare no competing financial interests.

---

**Figure 2–figure supplement 1.**
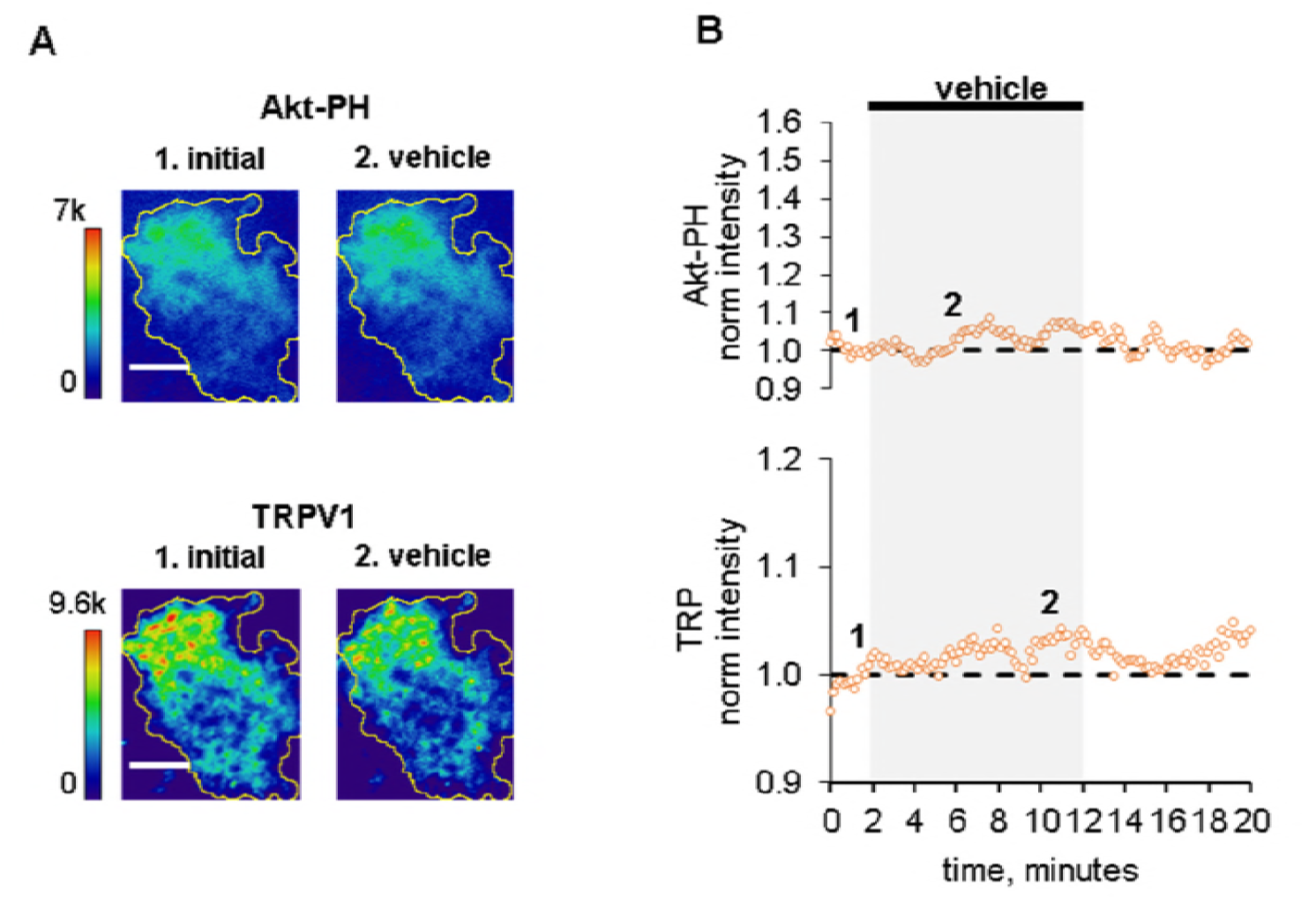
Vehicle does not increase PIP_3_ or recruit TRPV1 to PM. (A) TIRF images of a representative F-11 cell transfected with TrkA/p75_NTR_, TRPV1 and Akt-PH. Images labeled 1 were collected before vehicle application and those labeled 2 were collected at the time points labeled in B. Scale bar is 10 µm. LUT bars represent background-subtracted pixel intensities. The yellow border represents the outline of the cell footprint. (Top) Fluorescence intensity from Akt-PH. (Bottom) Fluorescence intensity from TRPV1. (B) Time course of vehicle-induced changes in fluorescence intensity for the cell shown in A. Vehicle was applied during the times indicated by the black bar/gray shading. Intensity at each time point was measured as the mean gray value within the footprint (yellow outline in A). Data were normalized to the mean intensity values during the two minutes prior to vehicle application.

**Figure 2–figure supplement 2.**
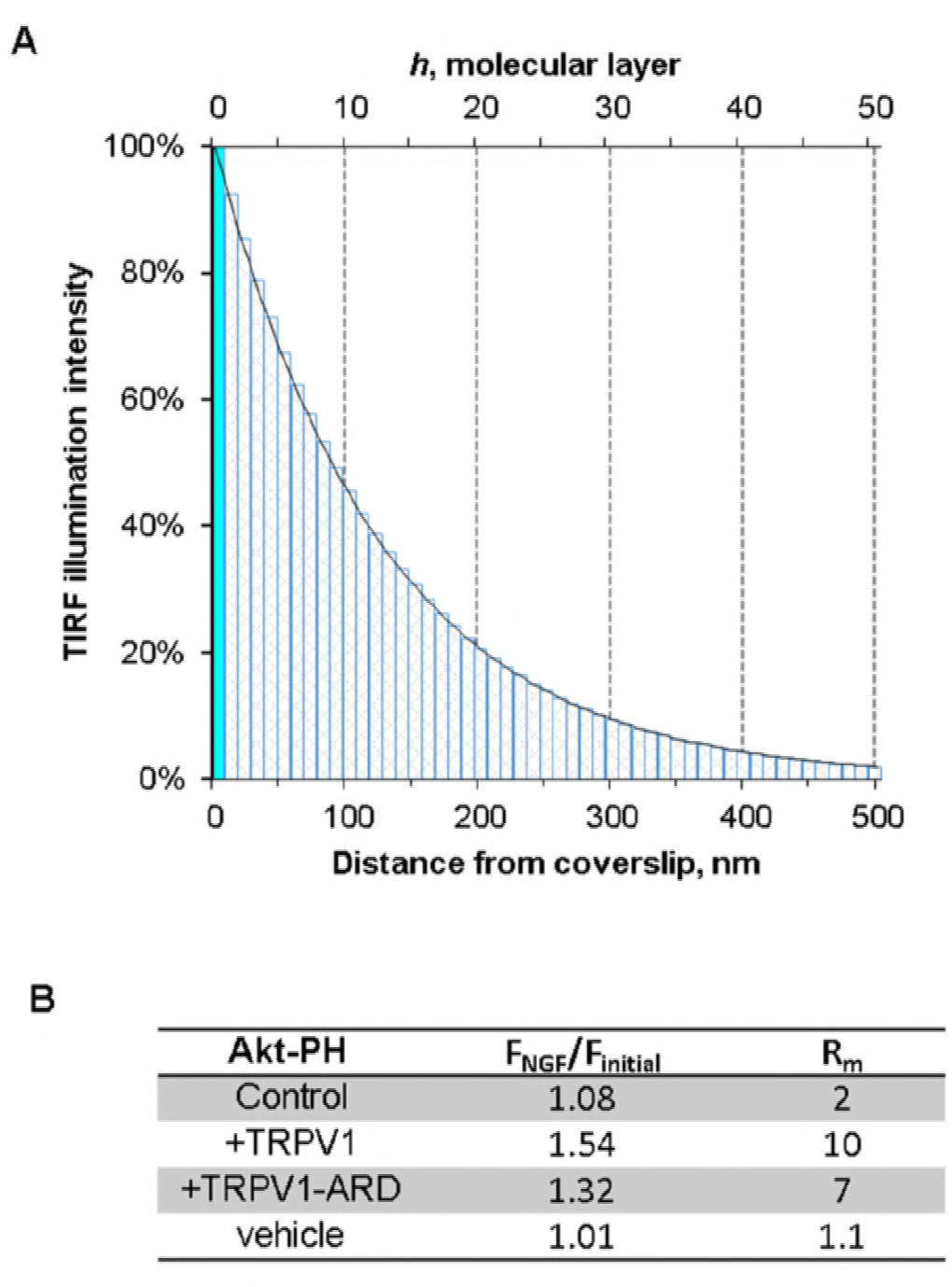
Model for TIRF illumination and estimation of Akt-PH translocation to the PM. (A) TIRF illumination intensity over distance in nanometers and molecular layers according to our model, see Materials and Methods. Refractive indexes of solution, coverslip and an incidence angle were all determined by our experimental conditions. Bars represent molecular layers, solid fill – membrane layer, pattern fill – cytosol layers. (B) Table for measured Akt-PH TIRF fluorescence (FNGF / Finitial), and estimated ratio of molecules at the membrane after NGF to that before NGF (Rm) Control is measurement of data in Figure 3A, TRPV1 – from Figure 3B, TRPV1-ARD–from Figure 5, vehicle – from Figure 2C. We consider the membrane and associated proteins to reside in layer h0. At rest, we assume that Akt-PH molecules are distributed evenly throughout layers h0-h49. We also assumed a fixed number of molecules in the field and that the only NGF-induced change was a redistribution of molecules among layers.

**Figure 3–figure supplement 1.**
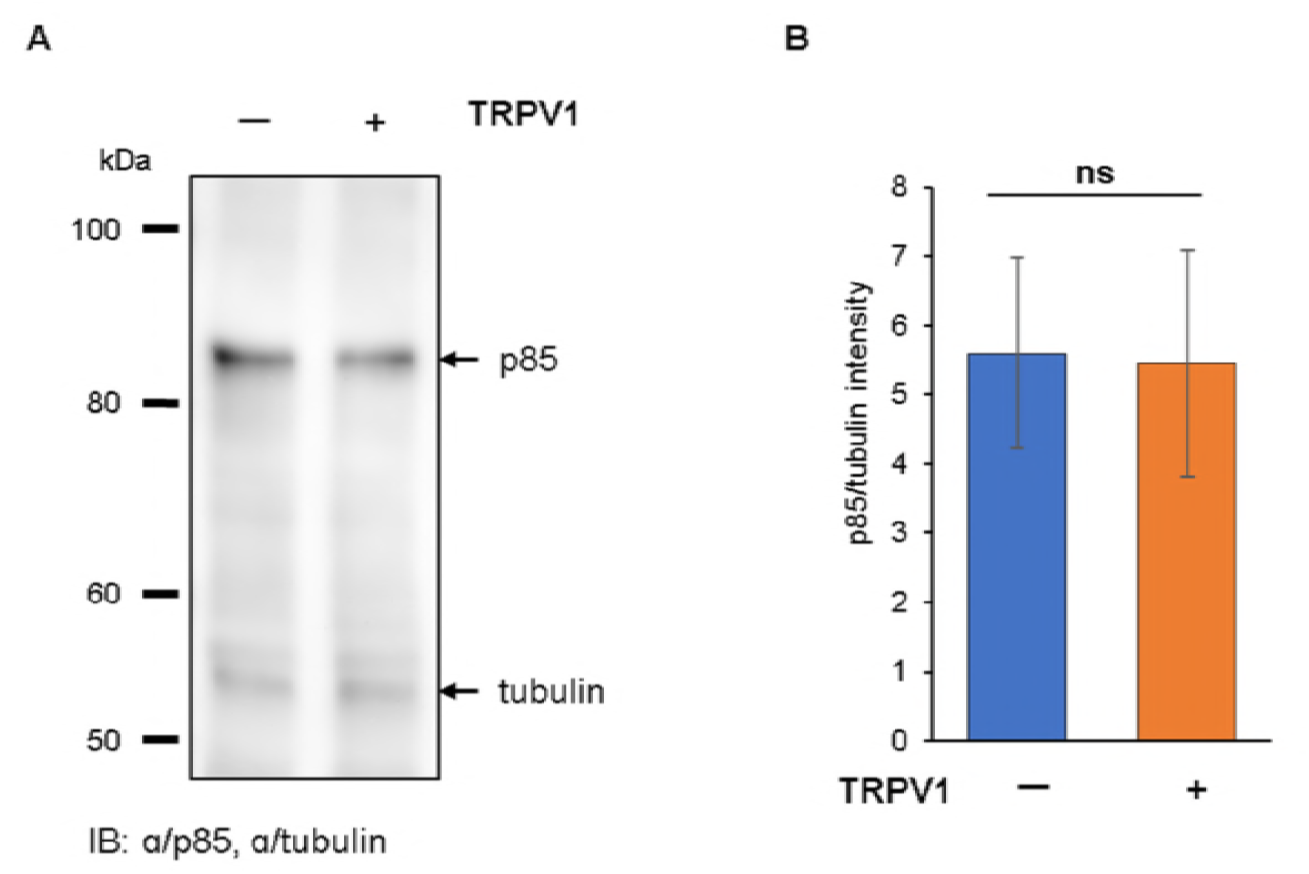
TRPV1 co-expression does not alter PI3K expression. (A) Representative immunoblot for p85 expression in F-11 cells expressing -/+ TRPV1. Membrane was stained for p85 and tubulin simultaneously. (B) Relative p85 expression in F-11 cells expressing -/+ TRPV1. Immunoblots were quantified as described in Methods (n = 5 independent experiments). Mean ± SEM pixel intensity are plotted normalized to the tubulin band on each blot. Student’s t-test two-tailed p=0.95.

**Figure 4–figure supplement 1.**
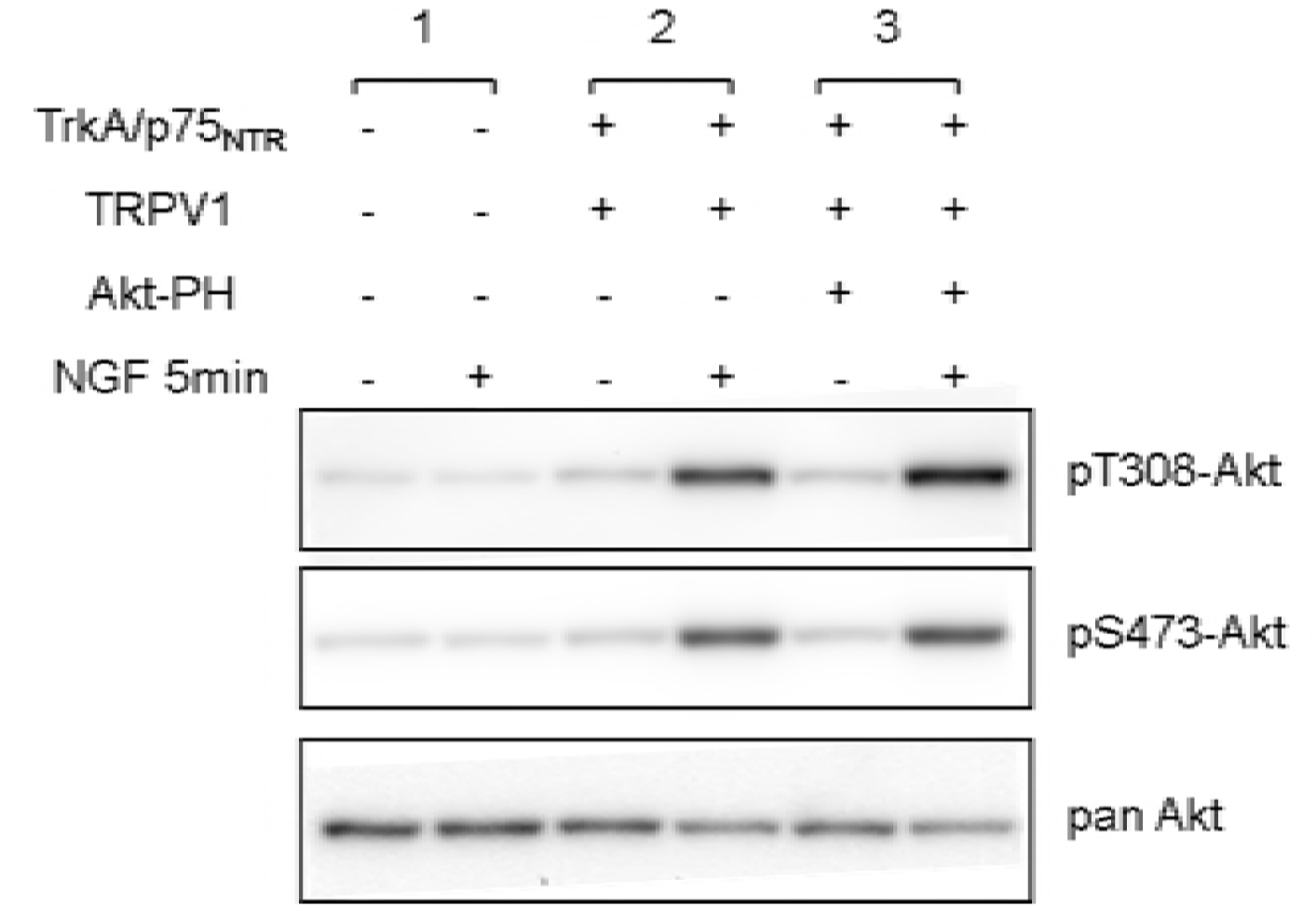
Akt-PH expression does not interfere with NGF-induced Akt phosphorylation. Immunoblot analysis for Akt-phosphorylation in cells lacking TrkA/p75_NTR_ (pair of lanes #1) treated with vehicle or NGF 100 ng/ml for 5 min. Cells lacking Akt-PH (pair of lanes #2) have comparable NGF-induced Akt-phosphorylation to cells transfected with Akt-PH (pair of lanes #3).

**Figure 4–figure supplement 2.**
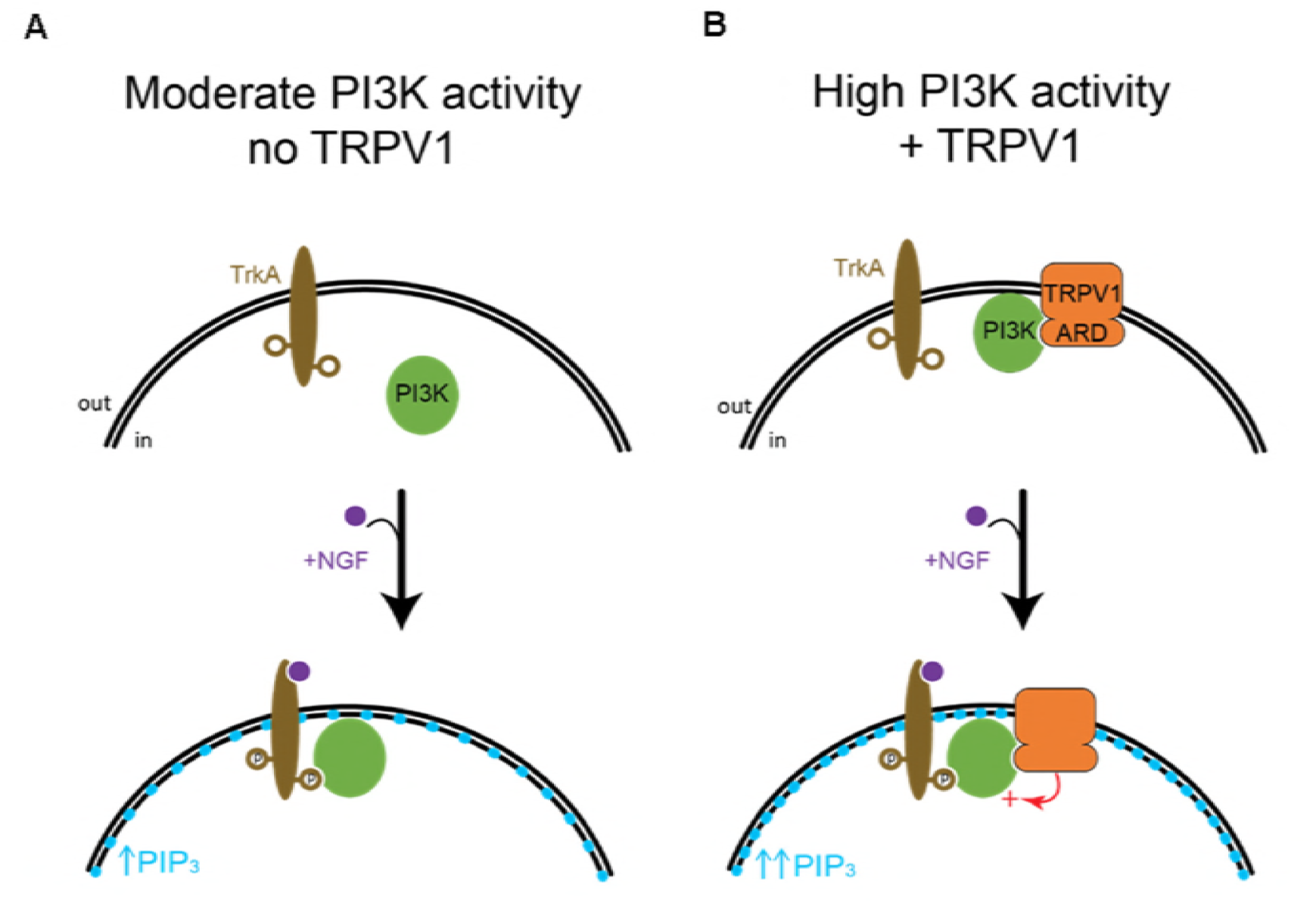
Simplified representation of PI3K activity. (A) In cells not expressing TRPV1, addition of NGF leads to moderate PI3K activity (represented as moderate increase in PIP_3_ levels, represented by blue circles in the inner membrane leaflet). (B) In the presence of TRPV1, addition of NGF leads to high PI3K activity (represented as high PIP_3_ levels). Potentiation is represented by the red arrow/+ symbols.

**Figure 5–figure supplement 1.**
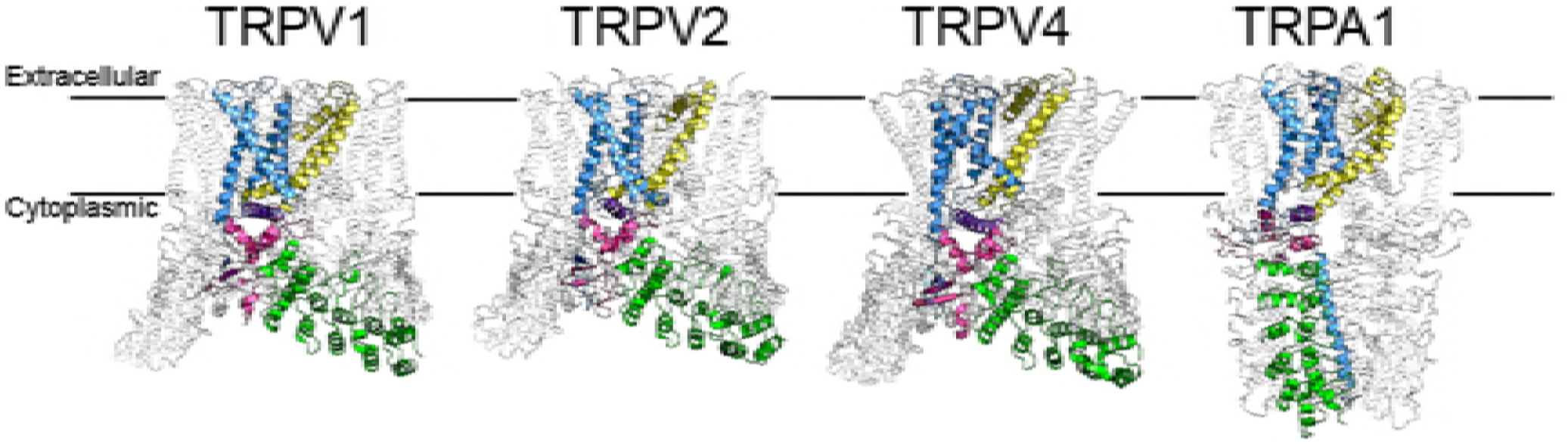
Comparison of TRP channels Cryo-EM structures. One subunit of each channel is colored as follows: ARD – green, TRP helix – purple, Pre-S1 – pink, Pore – yellow, all other parts of the subunit are depicted in blue. The remaining three subunits are outlined in gray for clarity. Cryo-EM structures of rat TRPV1, rat TRPV2, frog TRPV4 and human TRPA1 from PDB IDs 3J5P, 5HI9, 6BBJ, 3J9P, respectively, are shown.

**Figure 5–figure supplement 2.**
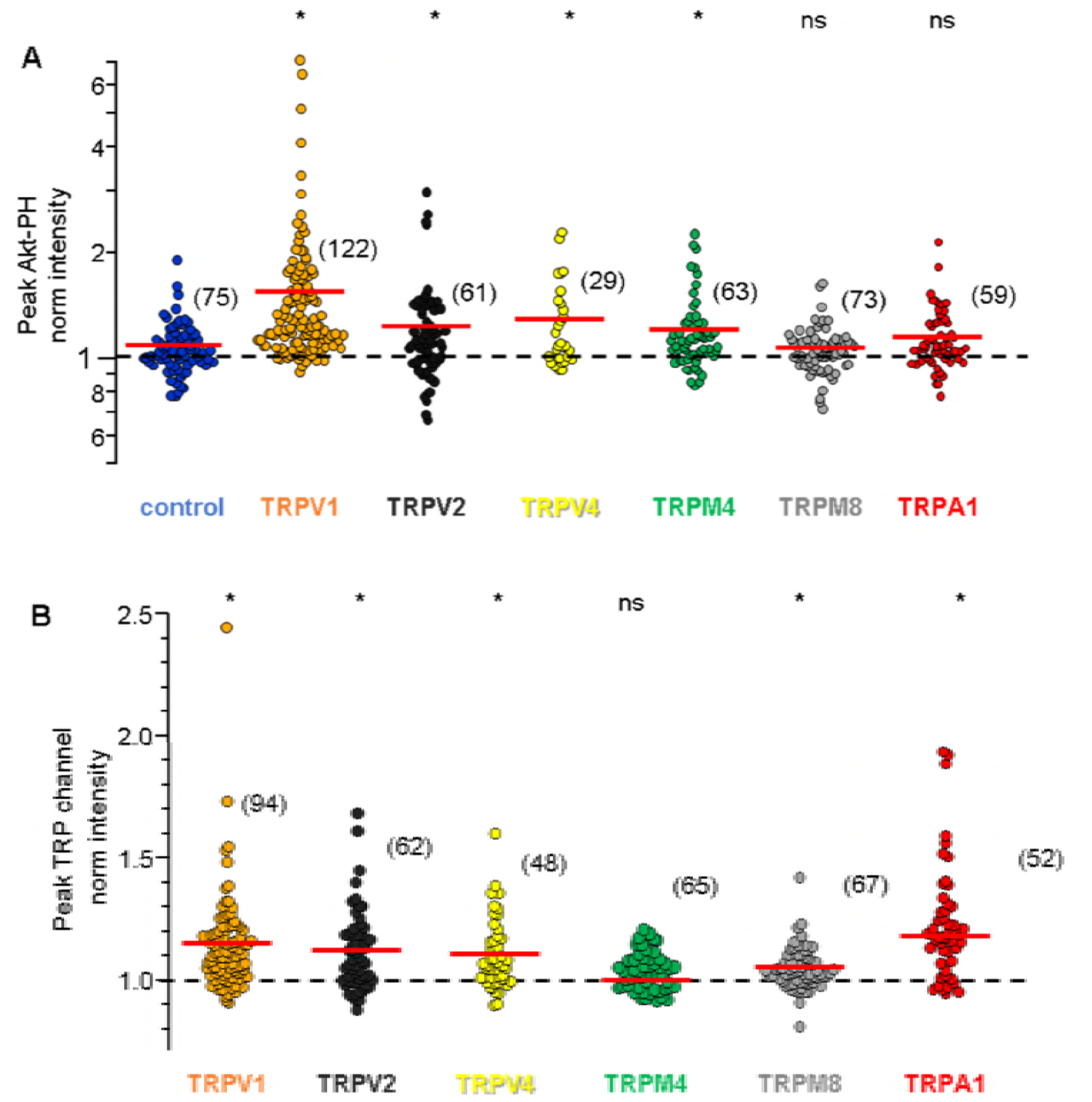
Potentiation of PI3K and NGF-induced trafficking are conserved among TRPV channels. NGF-induced change in fluorescence intensity. Each data point represents one cell. The red bars indicate the mean (see Table 1 for values). The y-axes are logarithmic. (A) Averaged normalized Akt-PH intensity during NGF application (6-8 minutes) from cells transfected with TrkA/p75_NTR_ and Akt-PH and: (A) no channel (control; blue; n=75); TRPV1 (orange; n=122); TRPV2 (black; n=61); TRPV4 (yellow; n=29); TRPM4 (green; n=63); TRPM8 (gray; n=73); TRPA1 (red; n=59). The y-axis is logarithmic. Asterisks represent statistical significance from Student’s t-test, comparing each TRP dataset to control with p<0.05, adjusted post-hoc to p<0.008 (values listed in Table 1). (B) Averaged normalized TRP channel intensity during NGF application (10-12 minutes). Color scheme as in (A) with the cell numbers as follows: (C) TRPV1 (n=94); TRPV2 (n=62); TRPV4 (n=48); (D) TRPM4 (n=65); TRPM8 (n=67); TRPA1 (n=52). Asterisks represent statistical significance from paired t-test comparing pre- to during-NGF values within each TRP dataset, with p values listed in Table 2.

**Figure 5–figure supplement 3.**
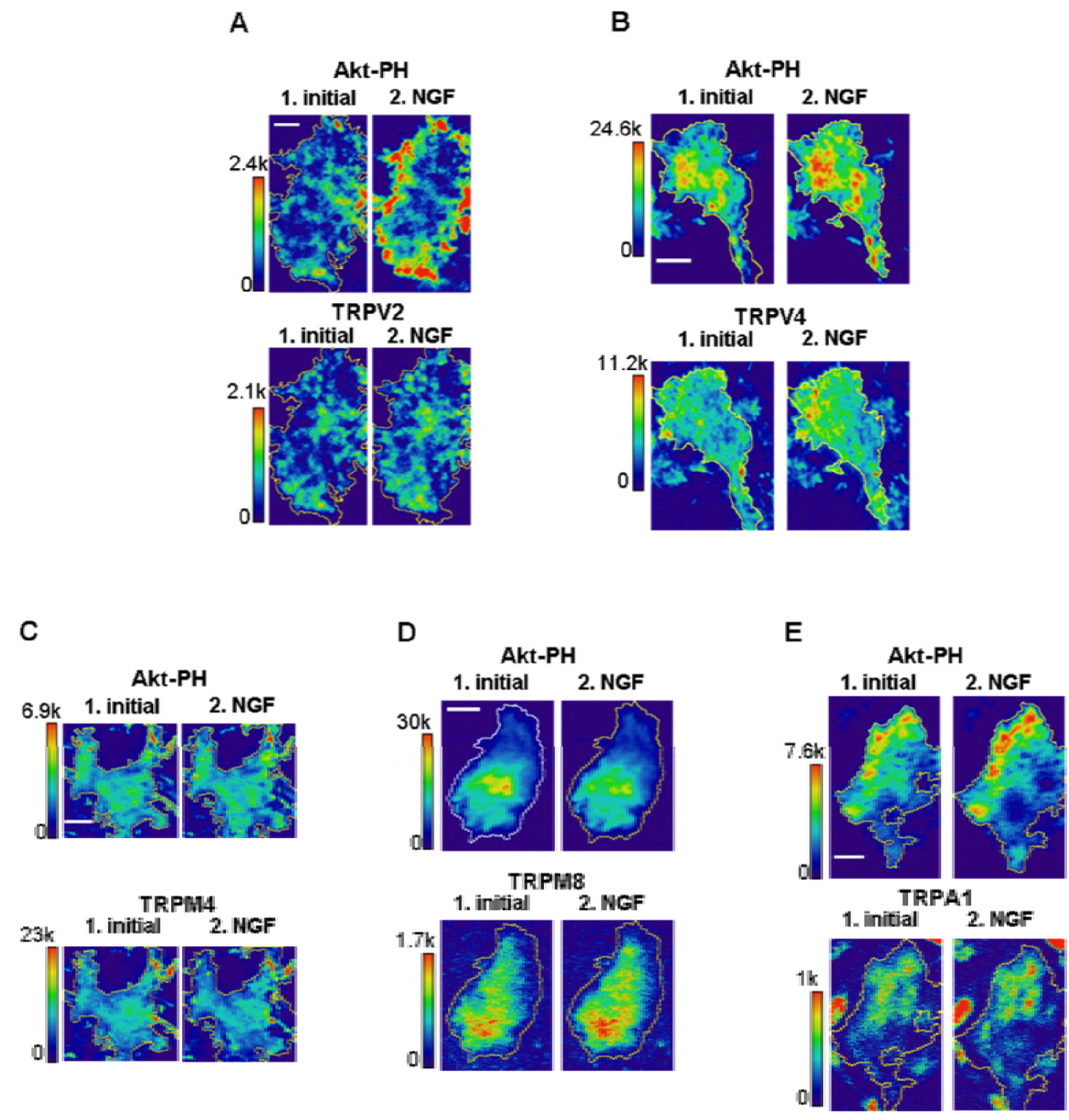
Representative images of NGF-induced recruitment Akt-PH and TRP channels to the PM. Representative images of F-11 cells transfected with TrkA/p75_NTR_, Akt-PH and one of the following: (A) TRPV2; (B) TRPV4; (C) TRPM4; (D) TRPM8; and (E) TRPA1. Timing of images and labels as in Figure 2A. Scale bar is 10 µm. LUT bar is background subtracted pixel intensities. Yellow outline represents the cell footprint.

